# Genetic meta-analysis identifies 9 novel loci and functional pathways for Alzheimer’s disease risk

**DOI:** 10.1101/258533

**Authors:** Iris E Jansen, Jeanne E Savage, Kyoko Watanabe, Julien Bryois, Dylan M Williams, Stacy Steinberg, Julia Sealock, Ida K Karlsson, Sara Hägg, Lavinia Athanasiu, Nicola Voyle, Petroula Proitsi, Aree Witoelar, Sven Stringer, Dag Aarsland, Ina S Almdahl, Fred Andersen, Sverre Bergh, Francesco Bettella, Sigurbjorn Bjornsson, Anne Brækhus, Geir Bråthen, Christiaan de Leeuw, Rahul S Desikan, Srdjan Djurovic, Logan Dumitrescu, Tormod Fladby, Timothy Homan, Palmi V Jonsson, Steven J Kiddle, K Arvid Rongve, Ingvild Saltvedt, Sigrid B. Sando, Geir Selbæk, Nathan Skenne, Jon Snaedal, Eystein Stordal, Ingun D. Ulstein, Yunpeng Wang, Linda R White, Jens Hjerling-Leffler, Patrick F Sullivan, Wiesje M van der Flier, Richard Dobson, Lea K. Davis, Hreinn Stefansson, Kari Stefansson, Nancy L Pedersen, Stephan Ripke, Ole A Andreassen, Danielle Posthuma

## Abstract

Late onset Alzheimer’s disease (AD) is the most common form of dementia with more than 35 million people affected worldwide, and no curative treatment available. AD is highly heritable and recent genome-wide meta-analyses have identified over 20 genomic loci associated with AD, yet only explaining a small proportion of the genetic variance indicating that undiscovered loci exist. Here, we performed the largest genome-wide association study of clinically diagnosed AD and AD-by-proxy (71,880 AD cases, 383,378 controls). AD-by-proxy status is based on parental AD diagnosis, and showed strong genetic correlation with AD (*r_g_*=0.81). Genetic meta analysis identified 29 risk loci, of which 9 are novel, and implicating 215 potential causative genes. Independent replication further supports these novel loci in AD. Associated genes are strongly expressed in immune-related tissues and cell types (spleen, liver and microglia). Furthermore, gene-set analyses indicate the genetic contribution of biological mechanisms involved in lipid-related processes and degradation of amyloid precursor proteins. We show strong genetic correlations with multiple health-related outcomes, and Mendelian randomisation results suggest a protective effect of cognitive ability on AD risk. These results are a step forward in identifying more of the genetic factors that contribute to AD risk and add novel insights into the neurobiology of AD to guide new drug development.

## Main text

Alzheimer′s disease (AD) is the most frequent neurodegenerative disease with roughly 35 million affected to date.^1^ Results from twin studies indicate that AD is highly heritable, with estimates ranging between 60 and 80%.^2^ Genetically, AD can be roughly divided into 2 subgroups: 1) familial early-onset cases that are relatively often explained by rare variants with a strong effect,^3^ and 2) late-onset cases that are influenced by multiple common variants with low effect sizes.^4^ Segregation analyses have linked several genes to the first subgroup, including *APP^5^, PSEN1^6^* and *PSEN2^7^.* The identification of these genes has resulted in valuable insights into a molecular mechanism with an important role in AD pathogenesis, the amyloidogenic pathway,^8^ providing a prominent example of how gene discovery can add to biological understanding of disease aetiology.

Besides the identification of a few rare genetic factors (e.g. *TREM2^9^* and *ABCA7^10^*), genome-wide association studies (GWAS) have mostly discovered common risk variants for the more complex late-onset type of AD. *APOE* is the strongest genetic risk locus for late-onset AD, where heterozygous and homozygous Apoe ε4 carriers are predisposed for a 3-fold and 15-fold increase in risk, respectively.^11^ A total of 19 additional GWAS loci have been described using a discovery sample of 17,008 AD cases and 37,154 controls, followed by replication of the implicated loci with 8,572 AD patients and 11,312 controls.^4^ The currently more than 20 confirmed AD risk loci explain only a fraction of the heritability of AD and increasing the sample size is likely to boost the power for detection of more common risk variants, which will aid in understanding biological mechanisms involved in the risk for AD.

In the current study, we included 455,258 individuals of European ancestry, meta analysed in 3 stages (**Figure 1**). These consisted of 24,087 clinically diagnosed late-onset AD cases, paired with 55,058 controls (phase 1). In phase 2, we analysed an AD-by-proxy phenotype, based on individuals in the UK Biobank (UKB) for whom parental AD status was available (N proxy cases=74,793; N proxy controls=328,320; **Online Methods**). The value of the usage of by-proxy phenotypes for GWAS was recently demonstrated by Liu et al^12^ for 12 common diseases. In particular for AD, Liu et al^12^ report substantial gains in statistical power by using a proxy phenotype, based on simulations and confirmed using empirical data from the 1^st^ release of the UKBiobank. We here apply the proxy phenotype strategy for AD in the UKBv2 sample. In this sample, parental diagnosis for AD was available for N=376,113 individuals, of whom 393 individuals had a known diagnosis of AD themselves (identified from medical register data). The high heritability of AD implies that case status for offspring can to some extent be inferred from parental case status and that offspring of AD parents are likely enriched for a higher genetic AD risk load. We thus defined individuals with one or two parents with AD as proxy cases (N=47,793), while putting more weight on the proxy cases with 2 parents. Similarly, the proxy controls include subjects with 2 parents without AD (N=328,320), where older cognitively normal parents were given more weight as proxy controls to account for the higher likelihood that younger parents may still develop AD. As the proxy phenotype is not a pure measure of an individual′s AD status and may include individuals that never develop AD, genetic effect sizes will be somewhat underestimated. However, the proxy case-control sample is very large (N proxy cases=47,793; N proxy controls=328,320), and therefore increases power to detect genetic effects for AD substantially.^12^ We first analysed the clinically defined case control samples separately from the by-proxy case control sample to allow investigation of overlap in genetic signals for these two measurements of AD risk. Finally in phase 3, we meta analysed all individuals of phase 1 and phase 2 together, and tested for replication in an independent sample.

**Figure 1.**
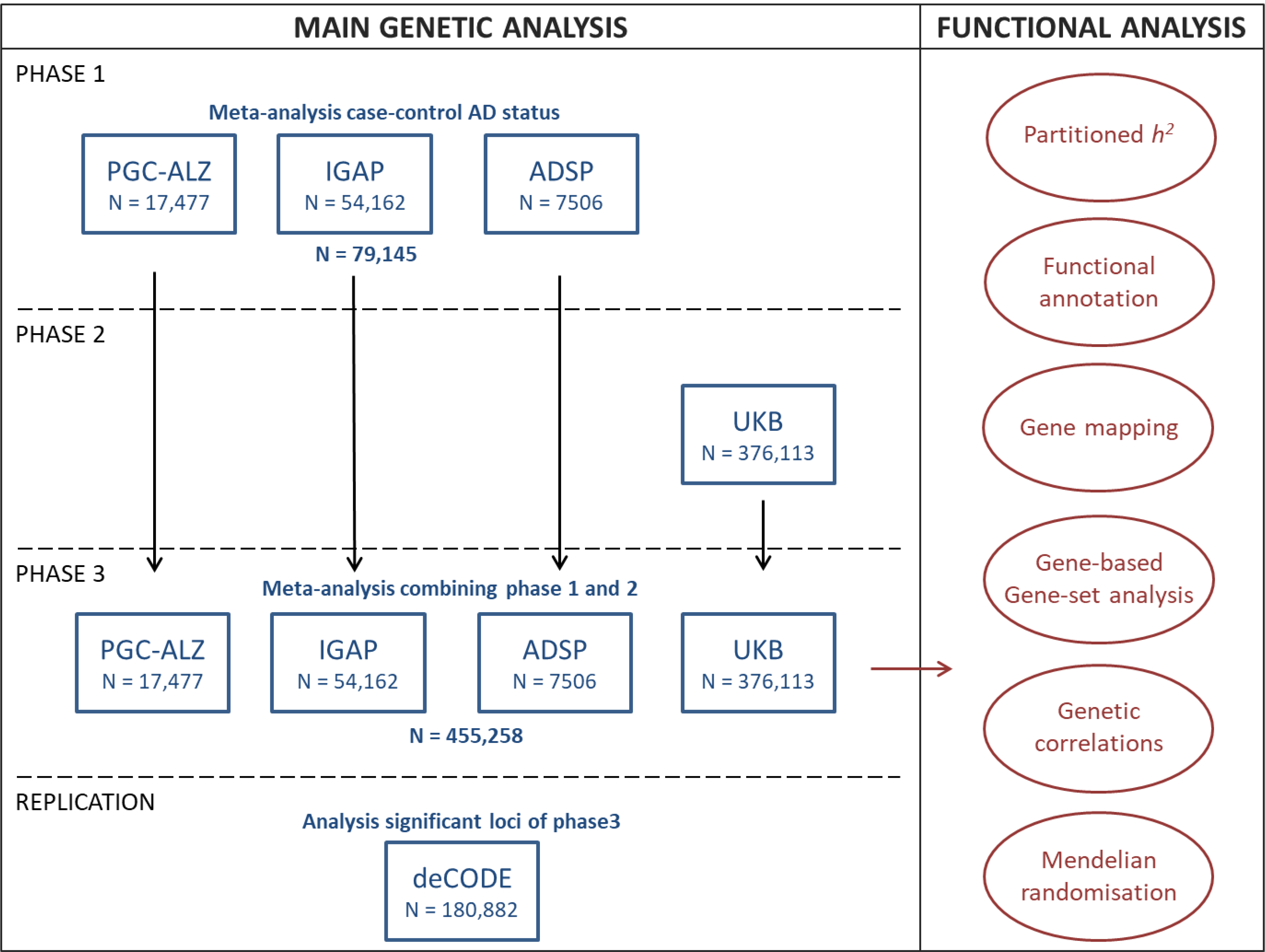
Overview of analyses steps. The main genetic analysis encompasses the procedures to detect GWAS risk loci for AD. The functional analysis part includes the *in silico* functional follow-up procedures with the aim to put the genetic findings in biological context. N = total of individuals within specified dataset.

### Genome-wide meta-analysis for AD status

Phase 1 involved a genome-wide meta-analysis for AD case-control status using cohorts collected as part of 3 independent main consortia (PGC-ALZ, IGAP and ADSP), totalling 79,145 individuals of European ancestry and 9,862,738 genetic variants passing quality control (**Figure 1, Supplementary Table 1**). The ADSP cohort obtained whole exome sequencing data from 4,343 cases and 3,163 controls, while the remaining datasets consisted of genotype single nucleotide polymorphism (SNP) array data. AD patients were diagnosed according to generally acknowledged diagnostic criteria, such as the NINCDS-ADRDA (See **Methods**). All cohorts for which we had access to the raw genotypic data were subjected to a standardized quality control pipeline, and GWA analyses were run per cohort and then included in a meta-analysis, alongside one dataset (IGAP) for which only summary statistics were available (see **Methods**). The full sample liability SNP-heritability (*h^2^_SNP_*), estimated with the more conservative LD Score regression (LDSC) method, was 0.055 (SE=0.0099), implying that 5.5% of AD heritability can be explained by the tested SNPs. This is in line with previous estimates for IGAP (6.8%) also estimated by LDSC regression method, which is based on summary statistics.^13,14^ We do note that previously reported estimates using a method based on raw genotypes (Genome-wide Complex Trait Analysis, GCTA), estimated that up to 53% of total phenotypic variance in AD could be explained by common SNPs, of which up to 6% could be explained by *APOE* alone, up to 13% by the then known variants, and up to 25% by undiscovered loci.^15,16^ The conservative LDSC estimate of *h^2^*_SNP_ is presumably a consequence of the underlying LDSC algorithm which is based on common HapMap SNPs and excludes all variants with extreme associations.

The **λ**_GC_=1.10 indicated the presence of inflated genetic signal compared to the null hypothesis of no association. The linkage disequilibrium (LD) score intercept^14^ was 1.044 (SE=0.0084) indicating that most inflation could be explained by polygenic signal (**Supplementary Figure 1**). In the meta-analysis of AD case-control status, 1,067 variants indexed by 51 lead SNPs in approximate linkage equilibrium (*r^2^*<0.1) reached genome-wide significance (GWS; P<5×10^−8^) (**Supplementary Figure 1; Supplementary Table 2**). These were located in 18 distinct genomic loci (**Table 1**). 15 of these loci confirmed previous findings (Lambert et al^4^) in a sample partially overlapping with that of the current study. The 3 remaining loci (lead SNPs* rs7657553, rs11257242 and rs2632516) have been linked more recently to AD in a genetic study^17^ of AD-related cholesterol levels while conditioning on lipid levels and in a transethnic genome-wide association study of AD.^18^

**Table 1.**
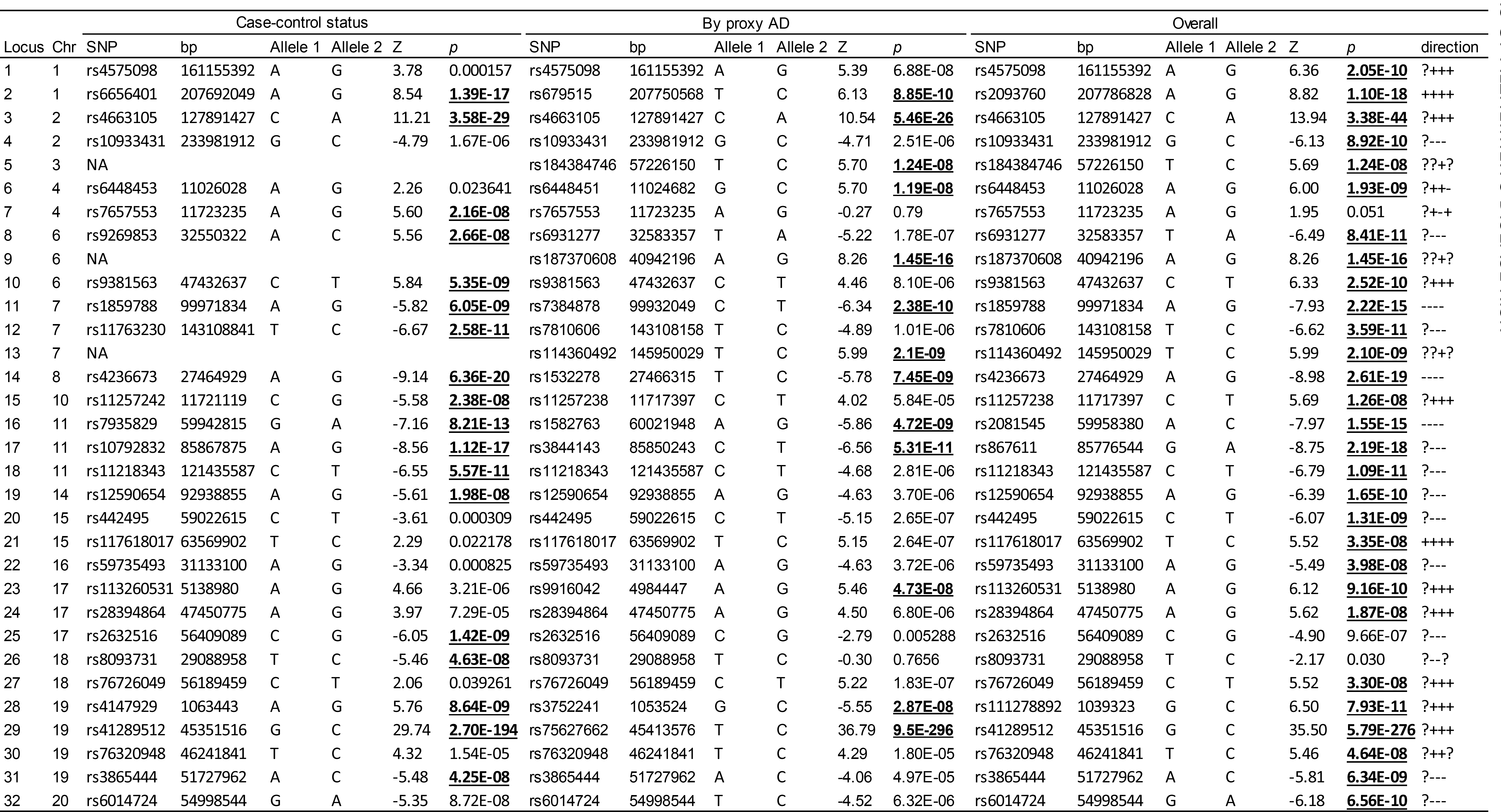
Summary statistics for the meta-analysis of case-control status, by proxy phenotype and both. Independent lead SNPs are defined by *r^2^* <. Indistinct genomic loci are >250kb apart. Alleles = the effect and non-effect allele. OR = odds ratio, only displayed for dichotomous phenotype, beta = effect size of the first allele displayed in the corresponding alleles column. Meta-analysis effect direction (column V) is in the following order: ADSP, IGAP, UKB, PGC ALZ, note that the first cohort is often missing as this concerns exome sequencing data. Corrected P value for significance = 5E-08 (marked as boldfaced and underlined values). Note that the lead SNP can differ between the distinct analyses, while it tags the same locus.

We next (phase 2) performed a GWAS for AD-by-proxy using 376,113 individuals of European ancestry from the UKB version 2 release using parental AD status weighted by age and corrected for population frequency to construct an AD-by-proxy status (**Figure 1; see Methods**). The LD score intercept was 1.022 (SE=0.0099) indicating that most of the inflation in genetic signal (**λ**_GC_=1.071) could be explained by polygenic signal (**Supplementary Figure 1B**). For AD-by-proxy, 719 GWS variants were indexed by 61 lead SNPs in approximate linkage equilibrium (*r^2^*<0.1) reached genome-wide significance (*P*<5×10^−8^), located in 13 loci (**Supplementary Figure 1A**). Of these, 8 loci overlapped with the significantly associated loci identified for clinical AD case control status (**Table 1**).

We observed a strong genetic correlation of 0.81 (SE= 0.185, using LDSC) between AD status and AD-by-proxy, indicating substantial overlap between genetic effects beyond shared GWS SNPs. Sign concordance tests indicated that 50.4% of all LD-independent (*r^2^* <0.1) genome wide SNPs (significant and non-significant) had consistent direction of effects between the two phenotypes (N=344,581 overlapping SNPs), slightly greater than the chance expectation of 50% (exact binomial test *P*=2.45×10^−7^). Of the 51 lead SNPs identified by the case-control meta-analysis, all were available in UKB and 96.1% were sign-concordant (*P*=2.98×10^−12^), while of the 61 GWS lead SNPs identified in UKB, 48 were available in the case control meta-analysis and 99.7% of these were sign-concordant (*P*=5.98×10^−14^). Such substantial overlap suggests that the AD-by-proxy phenotype captures a large part of the associated genetic effects on AD.

Given the high genetic overlap, in phase 3, we conducted a meta-analysis on the clinical AD case-control GWAS and the AD-by-proxy GWAS (**Figure 1**), comprising a total sample size of 455,258 (71,880 (proxy) cases and 383,378 (proxy) controls). The LD score intercept was 1.0018 (SE=0.0109) indicating again that most of the inflation in genetic signal (**λ**_GC_=1.0833) could be explained by polygenic signal (**Supplementary Figure 1b**). There were 2,357 GWS variants, which were represented by 94 lead SNPs, located in 29 loci (**Table 1**, **Figure 2**). These included 15 of the 18 loci detected in our case-control analyses, all of the 13 detected in the AD-by-proxy analyses, as well as 9 loci that were sub-threshold in both individual analyses but reached significance in the meta-analysis. All 2,160 GWS SNPs that were available in both the case control and AD-by-proxy sub-samples were sign concordant (exact binomial test *P*<1×10^−300^), including all of the 82 available independent lead SNPs (*P*=1.68×10^−23^). Association was found with both AD and AD-by-proxy for 22 (out of 27 overlapping) loci for which SNP(s) in each locus had a robust P-value (*P* < 0.05/94 independent signals). Of the 29 associated loci, 16 were previously identified by the GWAS of Lambert et al.,^4^ and 13 were not. Three of these (with lead SNPs rs184384746, rs187370608 and rs114360492) were only available in the UKB cohort (**Table 1**). Verifying our results against other ^9,19^ and more recent ^12,17,20^ genetic studies on AD, 4 loci (rs187370608, rs11257238, rs113260531 and rs28394864) were previously discovered, leaving 9 novel loci (rs4575098, rs184384746, rs6448453, rs114360492, rs442495, rs117618017, rs59735493, rs76726049 and rs76320948). Considering all loci of Lambert et al,^4^ we were unable to replicate 4 loci (*MEF2C, NME8, CELF1* and *FERMT2\**) at a GWS level (observed *P*-values were 1.6×10^−5^ to 0.0011), which was mostly caused by a lower association signal in the UKB dataset (**Supplementary Table 3**). By contrast, Lambert et al^4^ were unable to replicate the *DSG2* and *CD33* loci in the second stage of their study. In our study, *DSG2* is also not supported (meta-analysis *P*=0.030; UKB analysis *P*=0.766; **Table 1**), implying invalidation of this locus, while the *CD33* locus (rs3865444 in **Table 1**) is significantly associated with AD (meta analysis *P*=6.34 × 10^−9^; UKB analysis *P*=4.97 × 10^−5^), implying a genuine genetic association to AD risk.

**Figure 2.**
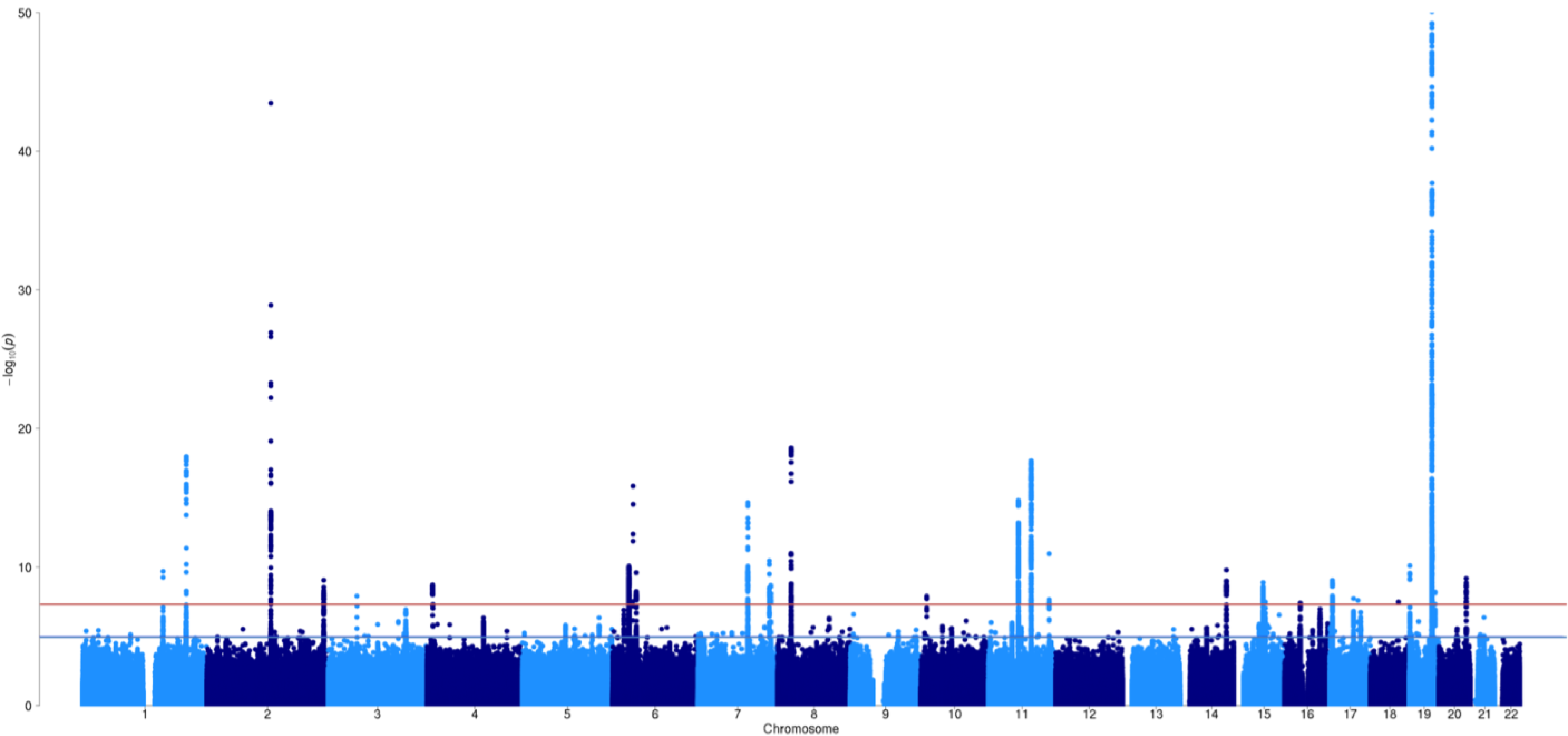
GWAS results for AD risk (N=455,258). Manhattan plot displays all associations per variant ordered according to their genomic position on the x-axis and showing the strength of the association with the −log10 transformed P-values on the y-axis. The y-axis is limited at 50 to enable visualization of *non-APOE* loci. The original −log10 for the APOE locus is 276.

Next, we aimed to find further support for the novel findings of the phase 3 meta analysis, by using an independent Icelandic cohort (deCODE^21,22^), including 6,593 AD cases and 174,289 controls (**Figure 1; see **Methods**; Supplementary Table 4**). We were unable to test two loci as the lead SNPs (and SNPs in high LD), either were not present in the 28,075 genomes of the Icelandic reference panel or were not imputed with sufficient quality. For 6 of the 7 novel loci tested for replication, we observed the same direction of effect in the deCODE cohort. Furthermore, 4 loci (rs6448453, rs442495, rs117618017, rs76320948) showed nominally significant association results (P<0.05) for the same SNP or a SNP in high LD (*r^2^* > 0.9) within the same locus (two-tailed binomial test *P*=1.9×10^−4^). The locus on chromosome 1 (rs45759098) was very close to significance (*P*=0.053). Apart from the novel loci, we also observed sign concordance for 95.6% of the lead SNPs in all loci from the meta-analysis (*P*=1.60×10^−20^) that were available in deCODE (out of 94). As an additional method of testing for replication using genome-wide polygenic score prediction,^23^ the current results explain 7.1% of the variance in clinical AD at a low best fitting *P*-threshold of 1.69×10^−5^ (*P*=1.80×10^−10^) in an independent sample of 761 individuals (see **Methods**). When excluding the *APOE-locus* (chr19: 45020859-45844508), the results explain 3.9% of the variance with a best fitting *P*-threshold of 3.5×10^−5^ (*P*=1.90×10^−6^).

### Functional interpretation of genetic variants contributing to AD and AD-by-proxy

Next, we conducted a number of *in silico* follow-up analyses to interpret our findings in a biological context. Functional annotation of all GWS SNPs (n=2,178) in the associated loci showed that SNPs were mostly located in intronic/intergenic areas, yet in regions that were enriched for chromatin states 4 and 5, implying effects on active transcription (**Figure 3A, 3B and 3C; Supplementary Table 5**). 24 GWS SNPs were exonic non-synonymous (ExNS) (**Figure 3A; Supplementary Table 6**) with likely deleterious implications on gene function. Converging evidence of strong association (Z> |7|) and a high observed probability of a deleterious variant effect (CADD^24^ score≥30) was found for rs75932628 (*TREM2*), rs142412517 (*TOMM40*) and rs7412 (*APOE).* The first two missense mutations are rare (MAF=0.002 and 0.001, respectively) and the alternative alleles were associated with higher risk for AD. The latter *APOE* missense mutation is the well-established protective allele Apoε2. The effect sizes for ExNS ranged from moderate to high. **Supplementary Tables 5 and 6** present a detailed annotation catalogue of variants in the associated genomic loci. Partitioned analysis,^25^ excluding SNPs with extremely large effect sizes (i.e. *APOE* variants) showed enrichment for *h^2^_SNP_* for variants located in H3K27ac marks (Enrichment=3.18, *P*=9.63×10^−5^), which are associated with activation of transcription, and in Super Enhancers (Enrichment=3.62, *P*=2.28×10^−4^), which are genomic regions where multiple epigenetic marks of active transcription are clustered (**Figure 3D; Supplementary Table 7**). Heritability was also enriched in variants on chromosome 17 (Enrichment=3.61, *P*=1.63×10^−4^) and we observed a trend of enrichment for variants with high minor allele frequencies (Enrichment=3.31, *P*=2.85×10^−3^), (**Supplementary Figure 3; Supplementary Tables 8 and 9**). Although a large proportion (23.9%) of the heritability can be explained by SNPs on chromosome 19, this enrichment is not significant, due to the large standard errors around this estimate (**Supplementary Table 8**). Overall these results suggest that, despite some nonsynonymous variants likely contributing to AD risk, most of the GWS SNPs are located in non-coding regions, and are enriched for regions that have an activating effect on transcription.

**Figure 3.**
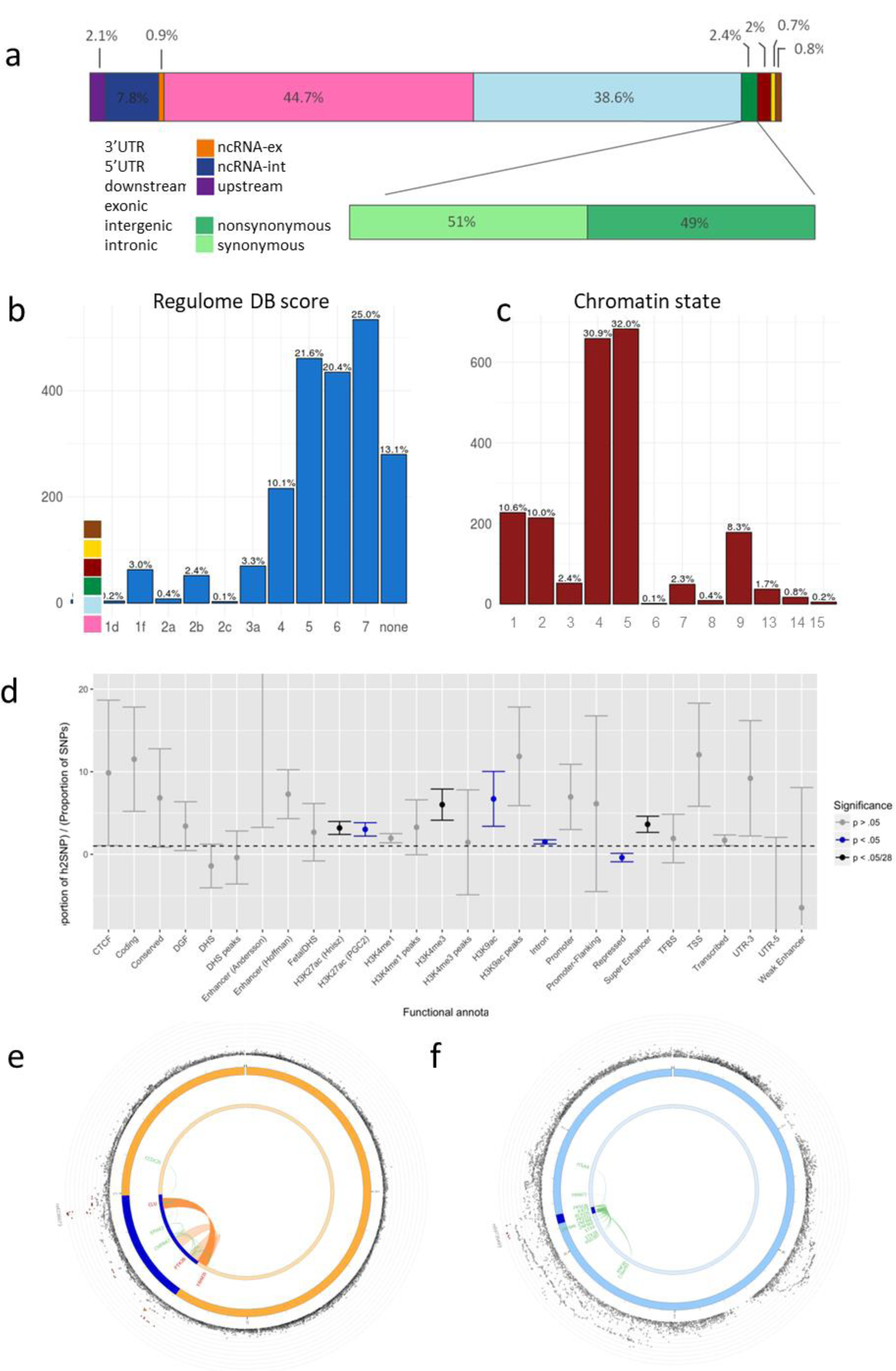
Functional annotation of association results. **a)** Heritability enrichment of 28 functional variant annotations calculated with stratified LD score regression. UTR=untranslated region; CTCF=CCCTC-binding factor; DHS=DNaseI hypersensitive site; TFBS=transcription factor binding site; DGF=DNAaseI digital genomic footprint; **b)** Functional effects of genome-wide significant variants in genomic risk loci of the meta-analysis the second bar shows the distribution for exonic variants only; **c)** Distribution of RegulomeDB score for variants in genomic risk loci, with a low score indicating a higher probability of having a regulatory function; **d)** Distribution of minimum chromatin state across 127 tissue and cell types for genome-wide significant variants in genomic risk loci, with lower states indicating higher accessibility and states 1-7 referring to open chromatin states. **e)** Zoomed-in circos plot of chromosome 8; **f)** Zoomed-in circos plot of chromosome 16. Circos plots show implicated genes by significant loci, where blue areas indicate genomic risk loci, green indicates eQTL associations and orange indicates chromatin interactions. Genes mapped by both eQTL and chromatin interactions are red. The outer layer shows a Manhattan plot containing the negative log10-transformed P-value of each SNP in the GWAS meta-analysis of AD. Full circos plots of all autosomal chromosomes are provided in Supplementary Figure 4.

### Implicated genes

To link the associated variants to genes, we applied three gene-mapping strategies implemented in FUMA^26^ (**Online Methods**). We used all SNPs with a P-value < 5×10^−8^ and *r^2^* of 0.6 with one of the independently associated SNPs, for gene-mapping. *Positional* gene-mapping aligned SNPs to 100 genes by their location within or immediately up/downstream (+/-10kb) of known gene boundaries, *eQTL (expression quantitative trait loci*) gene-mapping matched cis eQTL SNPs to 170 genes whose expression levels they influence in one or more tissues, and *chromatin interaction* mapping linked SNPs to 21 genes based on three-dimensional DNA-DNA interactions between each SNP’s genomic region and nearby or distant genes, which we limited to include only interactions between annotated enhancer and promotor regions (**Figure 3B and 3C; Supplementary Figure 4; Supplementary Tables 10 and 11**). This resulted in 192 uniquely mapped genes, 80 of which were implicated by at least two mapping strategies and 17 by all 3 (**Figure 4E**). Eight genes (*HLA-DRB5, HLA-DRB1, HLA-DQA, HLA-DQB1, KAT8, PRSS36, ZNF232* and *CEACAM19*) are particularly notable as they are implicated via eQTL association in the hippocampus, a brain region highly affected early in AD pathogenesis (**Supplementary Table 10**). Of special interest is the locus on chromosome 8 (rs4236673). In the GWAS by Lambert et al.^4^, this locus was defined as 2 distinct loci (*CLU* and *PTK2B*), while our meta-analysis specified this locus as a single locus based on LD-patterns. This is also supported by a chromatin interaction between the two regions (**Figure 3E**), which is observed in two immune-related tissues the spleen and liver (**Supplementary Table 11**). Chromosome 16 contains a locus implicated by long-range eQTL association (**Figure 3F**) clearly illustrating more distant genes can be affected by a genetic factor (**Figure 3F**) and emphasising the relevance of considering putative causal genes or regulatory elements not solely on the physical location but also on epigenetic influences. **Supplementary Figure 4** displays chromatin interactions for all chromosomes containing significant GWAS loci.

**Figure 4.**
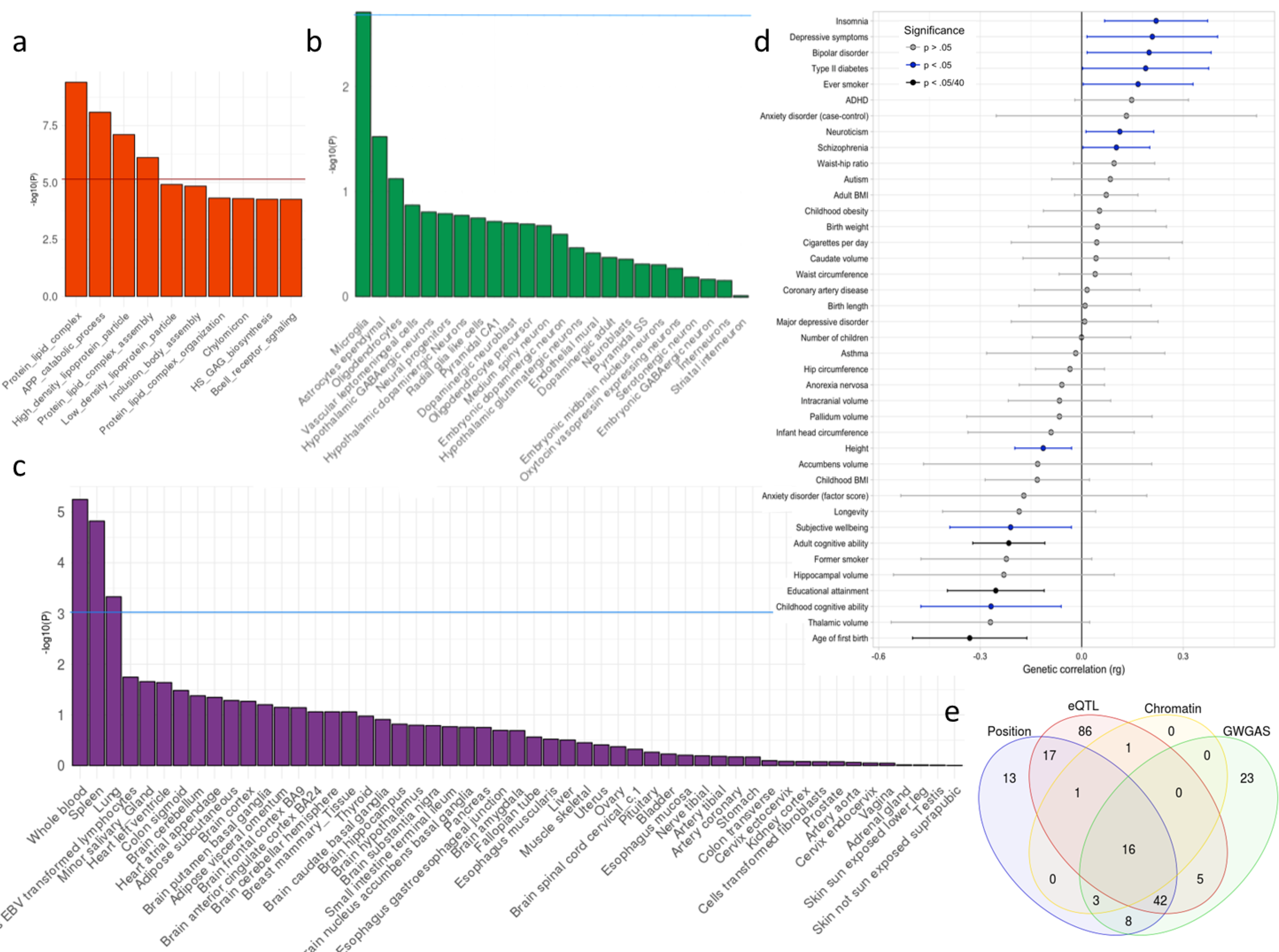
Functional implications based on gene-set analysis, genetic correlations and functional annotations. The gene-set results are displayed per category of biological mechanisms (A), brain cell types (B) and tissue types (C). The red horizontal lines indicates the significance threshold corrected for all gene-set tests of all categories, while the blue horizontal lines display the significance threshold corrected only for the number of tests within the three categories (i.e. gene-ontology, tissue expression, single cell expression). (D) Genetic correlations between AD and other heritable traits. (E) Venn diagram showing the number of genes mapped by four distinct strategies.

Although these gene-mapping strategies imply multiple putative causal genes per GWAS locus, several of these genes in the novel loci (and significantly replicated by the deCODE cohort) are of particular interest, as the genes have functional or previous genetic association to AD. For locus 1 in **Supplementary Table 10**, *ADAMTS4* encodes a protein of the ADAMTS family which has a function in neuroplasticity and has been extensively studied for their role in AD pathogenesis.^27^ For locus 19, the obvious most likely causal gene is *ADAM10,* as this gene has been associated with AD by research focusing on rare coding variants in ADAM10.^28^ However this is the first time that this gene is implicated as a common risk factor for AD. The lead SNP for locus 20 is a nonsynonymous variant in exon 1 of *APH1B,* which encodes for a protein subunit of the *γ*-secretase complex cleaving *APP.^29^* Although previously reported functional information on genes can be of great value, it is preferable to consider all implicated genes as putative causal factors to guide potential functional follow-up experiments.

We next performed genome-wide gene-based association analysis (GWGAS) using MAGMA.^30^ This method annotates SNPs to known protein-coding genes to estimate aggregate associations based on all SNPs in a gene. It differs from the gene-mapping strategies in FUMA as it provides a statistical gene-based test, whereas FUMA maps individually significant SNPs to genes. With GWGAS, we identified 97 genes that were significantly associated to AD (**Supplementary Figure 5; Supplementary Table 12**), of which 74 were also mapped by FUMA (**Figure 4E**). In total, 16 genes were implicated by all four strategies (**Supplementary Table 13**), of which 7 genes (*HLA-DRA, HLA-DRB1, PTK2B, CLU, MS4A3, SCIMP* and *RABEP1*) are not located in the APOE-locus, and therefore of high interest for further investigation.

### Gene-sets implicated in AD and AD-by-proxy

Using the gene-based P-values, we performed gene-set analysis for 6,994 biological-pathway based gene-sets, 53 tissue expression-based gene-sets and 39 brain single-cell expression based gene-sets (24 derived from mouse data and 15 derived from human data). We found four Gene Ontology^19^ gene-sets that were significantly associated with AD risk: *Protein lipid complex* (*P*=3.93×10^−10^), *Regulation of amyloid precursor protein catabolic process* (*P*=8.16×10^−09^), *High density lipoprotein particle* (*P*=7.81×10^−8^), and *Protein lipid complex assembly* (*P*=7.96×10^−7^) (**Figure 4A; Supplementary Tables 14 and 15**). Conditional analysis on the *APOE* locus showed associations with AD for these four gene-sets independent of the effect of *APOE,* as they remained significantly associated (*P*<0.0125), yet less strongly, suggesting that *APOE* is contributing a substantial part to the association signal, but does not completely drive the signal. There was overlap between genes included in the four gene-sets, and conditioning on each significant gene-set association showed that three gene-sets were associated with AD independently of each other (**Supplementary Tables 14 and 15**). All 25 genes of the *High density lipoprotein particle* pathway are also part of the *Protein lipid complex* (conditional analysis *P*=0.18), and these pathways are therefore not interpretable as independent associations.

Linking gene-based *P*-values to tissue and cell-type-specific gene-sets, no association survived the stringent Bonferroni correction, which corrected for all tested gene-sets (i.e. 6,994 GO categories, 54 tissues and 39 cell types). However, we did observe associations when correcting only for the number of tests within all tissue types or cell-types. This was the case for gene expression across immune-related tissues (**Figure 4C; Supplementary Table 16**), particularly whole blood (*P*=5.61×10^−6^), spleen (*P*=1.50×10^−5^) and lung (*P*=4.67×10^−4^). In brain single-cell expression gene-set analyses, we found associations for microglia, both in the mouse-based expression dataset (*P*=1.96×10^−3)^ (**Figure 4B; Supplementary Table 17**) and the human-based expression dataset (*P*=2.56×10^−3^) (**Supplementary Figure 6; Supplementary Table 18**).

### Cross-trait genetic influences

For a more comprehensive understanding of the genetic background of AD, we next tested whether AD is likely to share genetic factors with other phenotypes. This might reveal some functional insights about the genetic aetiology of AD. We conducted bivariate LD score^14^ regression to test for genetic correlations between AD and 41 other traits for which large GWAS summary statistics were available. We observed significant negative genetic correlations with adult cognitive ability (*r_g_*=-0.22, *P*=7.28×10^−5^), age of first birth (*r_g_*=-0.33, *P*=1.22×10^−4^), educational attainment (*r_g_*=-0.25, *P*=5.01×10^−4^), and confirmed a very strong positive correlation with previous GWAS of Alzheimer’s disease (*r_g_*=0.90, *P*=3.29×10^−16^) (**Figure 4D; Supplementary Table 19**).

We then used Generalised Summary-statistic-based Mendelian Randomisation^31^ (GSMR; See **Methods**) to test for potential credible causal associations of genetically correlated outcomes which may directly influence the risk for AD. Due to the nature of AD being a late onset disorder and summary statistics for most other traits being obtained from younger samples, we do not report tests for the opposite direction of potential causality (i.e. we did not test for a causal effect of a late-onset disease on an early onset disease). In this set of analyses, SNPs from the summary statistic of genetically correlated phenotypes were used as instrumental variables to estimate the putative causal effect of these ″exposure″ phenotypes on AD risk by comparing the ratio of SNPs’ associations with each exposure to their associations with AD outcome (See **Methods**). Association statistics were standardized, such that the reported effects reflect the expected difference in odds ratio (OR) for AD as a function of every SD increase in the exposure phenotype. We observed a protective effect of cognitive ability (OR=0.89, 95% confidence interval[CI]: 0.85-0.92, *P*=5.07×10^−9^), educational attainment (OR=0.88, 95%CI: 0.81-0.94, *P*=3.94×10^−4^), and height (OR=0.96, 95%CI: 0.94-0.97, *P*=1.84×10^−8^) on risk for AD (**Supplementary Table 20**; **Supplementary Figure 7**). No substantial evidence of pleiotropy was observed between AD and these phenotypes, with <1% of overlapping SNPs being filtered as outliers (**Supplementary Figure 7**).

### Discussion

By using a non-conventional approach of including a by-proxy phenotype for AD to increase sample size, we have identified 9 novel loci and gained novel biological knowledge on AD aetiology. Both the high genetic correlation between the standard case-control status and the UKB by proxy phenotype (*r_g_*=0.81) and the high rate of novel loci replication in the independent deCODE cohort, suggest that this strategy is robust. Through extensive in silico functional follow-up analysis, and in line with previous research,^20,32^ we emphasise the crucial causal role of the immune system rather - than immune response as a consequence of disease pathology - by establishing variant enrichments for immune-related body tissues (whole blood, spleen, liver) and for the main immune cells of the brain (microglia). Furthermore, we observe informative eQTL associations and chromatin interactions within immune-related tissues for the identified genomic risk loci. Together with the AD-associated genetic effects on lipid metabolism in our study, these biological implications strengthen the hypothesis that AD pathogenesis involves an interplay between inflammation and lipids, as lipid changes might harm immune responses of microglia and astrocytes, and vascular health of the brain.^33^

In accordance with previous clinical research, our study suggests an important role for protective effects of several human traits on AD. As an example, cognitive reserve has been proposed as a protective mechanism in which the brain aims to control brain damage with prior existing cognitive processing strategies.^34^ Our findings imply that some component of the genetic factors for AD might affect cognitive reserve, rather than being involved in AD pathology-related damaging processes, influencing AD pathogenesis in an indirect way through cognitive reserve. Similarly, in a largescale community-based study it was observed that AD incidence rates declined over decades, which was specific for individuals with at minimum a high school diploma.^35^ Combined with our Mendelian randomization results for educational attainment, this suggests that the protective effect of educational attainment on AD is influenced by genetics.

The results of this study could furthermore serve as a valuable resource (e.g. Supplementary Tables 10 and 13) for selection of promising genes for functional follow-up experiments and identify targets for drug development. We anticipate that functional interpretation strategies and follow-up experiments will result in a comprehensive understanding of late-onset AD aetiology, which will serve as a solid foundation for future AD drug development and stratification approaches.

## URLs

http://ukbiobank.ac.uk

https://www.ncbi.nlm.nih.gov/gap

http://fuma.ctglab.nl

http://ctg.cncr.nl/software/magma

http://genome.sph.umich.edu/wiki/METALProgram

https://github.com/bulik/ldsc

http://ldsc.broadinstitute.org/

https://data.broadinstitute.org/alkesgroup/LDSCORE/

http://www.genecards.org

http://www.med.unc.edu/pgc/results-and-downloads

http://software.broadinstitute.org/gsea/msigdb/collections.jsp

https://www.ebi.ac.uk/gwas/

https://github.com/ivankosmos/RegionAnnotator

http://cnsgenomics.com/software/gsmr/

## Acknowledgments

This work was funded by The Netherlands Organization for Scientific Research (NWO VICI 453-14-005) and the Sophia Foundation for Scientific Research (grant nr: S14-27). The analyses were carried out on the Genetic Cluster Computer, which is financed by the Netherlands Scientific Organization (NWO: 480-05-003), by the VU University, Amsterdam, The Netherlands, and by the Dutch Brain Foundation, and is hosted by the Dutch National Computing and Networking Services SurfSARA. The work was also funded by The Research Council of Norway (#251134, #248778, #223273, #213837, #225989), KG Jebsen Stiftelsen, The Norwegian Health Association, European Community’s JPND Program, ApGeM RCN #237250, and the European Community′s grant # PIAPP-GA-2011-286213 PsychDPC. This research has been conducted using the UK Biobank resource under application number 16406 and the public ADSP dataset, obtained through the Database of Genotypes and Phenotypes (dbGaP) under accession number phs000572 (see sections below).

Genotyping for the Swedish Twin Studies of Aging was supported by NIH/NIA grant R01 AG037985. Genotyping in TwinGene was supported by NIH/NIDDK U01 DK066134. WvdF is recipient of Joint Programming for Neurodegenerative Diseases (JPND) grants PERADES (ANR 13-JPRF-0001) and EADB (733051061). AddNeuroMed consortium was led by Simon Lovestone, Bruno Vellas, Patrizia Mecocci, Magda Tsolaki, Iwona Kłoszewska, Hilkka Soininen. This work was supported by InnoMed (Innovative Medicines in Europe), an integrated project funded by the European Union of the Sixth Framework program priority (FP6-2004 LIFESCIHEALTH-5). JB was supported by a grant from the Swiss National Science Foundation. JHL was supported by the Swedish Research Council (Vetenskapsrådet, award 2014-3863), the Wellcome Trust (108726/Z/15/Z), and the Swedish Brain Foundation (Hjärnfonden). NS was supported by the Wellcome Trust (108726/Z/15/Z). RD was supported by National Institute for Health Research University College London Hospital′s Biomedical Research Centre, Arthritis Research UK, the British Heart Foundation, Cancer Research UK, the Chief Scientist Office, the Economic and Social Research Council, the Engineering and Physical Sciences Research Council, the National Institute for Social Care and Health Research, and the Wellcome Trust (grant number MR/K006584/1), Innovative Medicines Initiative Joint Undertaking under EMIF grant agreement number 115372, resources of which are composed of financial contribution from the European Union′s Seventh Framework Program (FP7/2007-2013) and EFPIA companies′ in kind contribution. SJK was supported by an MRC Career Development Award in Biostatistics (MR/L011859/1).

We thank the International Genomics of Alzheimer′s Project (IGAP) for providing summary results data for these analyses. The investigators within IGAP contributed to the design and implementation of IGAP and/or provided data but did not participate in analysis or writing of this report. IGAP was made possible by the generous participation of the control subjects, the patients, and their families. The i-Select chips was funded by the French National Foundation on Alzheimer′s disease and related disorderacknows. EADI was supported by the LABEX (laboratory of excellence program investment for the future) DISTALZ grant, Inserm, Institut Pasteur de Lille, Université de Lille 2 and the Lille University Hospital. GERAD was supported by the Medical Research Council (Grant n° 503480), Alzheimer′s Research UK (Grant n° 503176), the Wellcome Trust (Grant n° 082604/2/07/Z) and German Federal Ministry of Education and Research (BMBF): Competence Network Dementia (CND) grant n° 01GI0102, 01GI0711, 01GI0420. CHARGE was partly supported by the NIH/NIA grant R01 AG033193 and the NIA AG081220 and AGES contract N01-AG-12100, the NHLBI grant R01 HL105756, the Icelandic Heart Association, and the Erasmus Medical Center and Erasmus University. ADGC was supported by the NIH/NIA grants: U01 AG032984, U24 AG021886, U01 AG016976, and the Alzheimer′s Association grant ADGC-10-196728. This paper represents independent research funded by the National Institute for Health Research (NIHR) Biomedical Research Centre at South London and Maudsley NHS Foundation Trust and King′s College London. The views expressed are those of the author(s) and not necessarily those of the NHS, the NIHR or the Department of Health.

The Alzheimer′s Disease Sequencing Project (ADSP) is comprised of two Alzheimer′s Disease (AD) genetics consortia and three National Human Genome Research Institute (NHGRI) funded Large Scale Sequencing and Analysis Centers (LSAC). The two AD genetics consortia are the Alzheimer′s Disease Genetics Consortium (ADGC) funded by NIA (U01 AG032984), and the Cohorts for Heart and Aging Research in Genomic Epidemiology (CHARGE) funded by NIA (R01 AG033193), the National Heart, Lung, and Blood Institute (NHLBI), other National Institute of Health (NIH) institutes and other foreign governmental and non-governmental organizations. The Discovery Phase analysis of sequence data is supported through UF1AG047133 (to Drs. Schellenberg, Farrer, Pericak-Vance, Mayeux, and Haines); U01AG049505 to Dr. Seshadri; U01AG049506 to Dr. Boerwinkle; U01AG049507 to Dr. Wijsman; and U01AG049508 to Dr. Goate and the Discovery Extension Phase analysis is supported through U01AG052411 to Dr. Goate, U01AG052410 to Dr. Pericak-Vance and U01 AG052409 to Drs. Seshadri and Fornage. Data generation and harmonization in the Follow-up Phases is supported by U54AG052427 (to Drs. Schellenberg and Wang). The ADGC cohorts include: Adult Changes in Thought (ACT), the Alzheimer′s Disease Centers (ADC), the Chicago Health and Aging Project (CHAP), the Memory and Aging Project (MAP), Mayo Clinic (MAYO), Mayo Parkinson′s Disease controls, University of Miami, the Multi-Institutional Research in Alzheimer′s Genetic Epidemiology Study (MIRAGE), the National Cell Repository for Alzheimer′s Disease (NCRAD), the National Institute on Aging Late Onset Alzheimer′s Disease Family Study (NIA-LOAD), the Religious Orders Study (ROS), the Texas Alzheimer′s Research and Care Consortium (TARC), Vanderbilt University/Case Western Reserve University (VAN/CWRU), the Washington Heights-Inwood Columbia Aging Project (WHICAP) and the Washington University Sequencing Project (WUSP), the Columbia University Hispanic Estudio Familiar de Influencia Genetica de Alzheimer (EFIGA), the University of Toronto (UT), and Genetic Differences (GD). The CHARGE cohorts are supported in part by National Heart, Lung, and Blood Institute (NHLBI) infrastructure grant HL105756 (Psaty), RC2HL102419 (Boerwinkle) and the neurology working group is supported by the National Institute on Aging (NIA) R01 grant AG033193. The CHARGE cohorts participating in the ADSP include the following: Austrian Stroke Prevention Study (ASPS), ASPS-Family study, and the Prospective Dementia Registry-Austria (ASPS/PRODEM-Aus), the Atherosclerosis Risk in Communities (ARIC) Study, the Cardiovascular Health Study (CHS), the Erasmus Rucphen Family Study (ERF), the Framingham Heart Study (FHS), and the Rotterdam Study (RS). ASPS is funded by the Austrian Science Fond (FWF) grant number P20545-P05 and P13180 and the Medical University of Graz. The ASPS-Fam is funded by the Austrian Science Fund (FWF) project I904), the EU Joint Programme Neurodegenerative Disease Research (JPND) in frame of the BRIDGET project (Austria, Ministry of Science) and the Medical University of Graz and the Steiermärkische Krankenanstalten Gesellschaft. PRODEM-Austria is supported by the Austrian Research Promotion agency (FFG) (Project No. 827462) and by the Austrian National Bank (Anniversary Fund, project 15435. ARIC research is carried out as a collaborative study supported by NHLBI contracts (HHSN268201100005C, HHSN268201100006C, HHSN268201100007C, HHSN268201100008C, HHSN268201100009C, HHSN268201100010C, HHSN268201100011C, and HHSN268201100012C). Neurocognitive data in ARIC is collected by U01 2U01HL096812, 2U01HL096814, 2U01HL096899, 2U01HL096902, 2U01HL096917 from the NIH (NHLBI, NINDS, NIA and NIDCD), and with previous brain MRI examinations funded by R01 HL70825 from the NHLBI. CHS research was supported by contracts HHSN268201200036C, HHSN268200800007C, N01HC55222, N01HC85079, N01HC85080, N01HC85081, N01HC85082, N01HC85083, N01HC85086, and grants U01HL080295 and U01HL130114 from the NHLBI with additional contribution from the National Institute of Neurological Disorders and Stroke (NINDS). Additional support was provided by R01AG023629, R01AG15928, and R01AG20098 from the NIA. FHS research is supported by NHLBI contracts N01-HC-25195 and HHSN268201500001I. This study was also supported by additional grants from the NIA (R01s AG054076, AG049607 and AG033040 and NINDS (R01 NS017950). The ERF study as a part of EUROSPAN (European Special Populations Research Network) was supported by European Commission FP6 STRP grant number 018947 (LSHG-CT-2006-01947) and also received funding from the European Community′s Seventh Framework Programme (FP7/2007-2013)/grant agreement HEALTH-F4-2007-201413 by the European Commission under the programme "Quality of Life and Management of the Living Resources" of 5th Framework Programme (no. QLG2-CT-2002-01254). High-throughput analysis of the ERF data was supported by a joint grant from the Netherlands Organization for Scientific Research and the Russian Foundation for Basic Research (NWO-RFBR 047.017.043). The Rotterdam Study is funded by Erasmus Medical Center and Erasmus University, Rotterdam, the Netherlands Organization for Health Research and Development (ZonMw), the Research Institute for Diseases in the Elderly (RIDE), the Ministry of Education, Culture and Science, the Ministry for Health, Welfare and Sports, the European Commission (DG XII), and the municipality of Rotterdam. Genetic data sets are also supported by the Netherlands Organization of Scientific Research NWO Investments (175.010.2005.011, 911-03-012), the Genetic Laboratory of the Department of Internal Medicine, Erasmus MC, the Research Institute for Diseases in the Elderly (014-93-015; RIDE2), and the Netherlands Genomics Initiative (NGI)/Netherlands Organization for Scientific Research (NWO) Netherlands Consortium for Healthy Aging (NCHA), project 050-060-810. All studies are grateful to their participants, faculty and staff. The content of these manuscripts is solely the responsibility of the authors and does not necessarily represent the official views of the National Institutes of Health or the U.S. Department of Health and Human Services. The three LSACs are: the Human Genome Sequencing Center at the Baylor College of Medicine (U54 HG003273), the Broad Institute Genome Center (U54HG003067), and the Washington University Genome Institute (U54HG003079). Biological samples and associated phenotypic data used in primary data analyses were stored at Study Investigators institutions, and at the National Cell Repository for Alzheimer′s Disease (NCRAD, U24AG021886) at Indiana University funded by NIA. Associated Phenotypic Data used in primary and secondary data analyses were provided by Study Investigators, the NIA funded Alzheimer′s Disease Centers (ADCs), and the National Alzheimer′s Coordinating Center (NACC, U01AG016976) and the National Institute on Aging Genetics of Alzheimer′s Disease Data Storage Site (NIAGADS, U24AG041689) at the University of Pennsylvania, funded by NIA, and at the Database for Genotypes and Phenotypes (dbGaP) funded by NIH. This research was supported in part by the Intramural Research Program of the National Institutes of health, National Library of Medicine. Contributors to the Genetic Analysis Data included Study Investigators on projects that were individually funded by NIA, and other NIH institutes, and by private U.S. organizations, or foreign governmental or nongovernmental organizations.

We thank the numerous participants, researchers, and staff from many studies who collected and contributed to the data. Summary statistics will be made available for download upon publication from http://ctglab.vu.nl.

## Author Contributions

I.E.J. and J.E.S. performed the analyses. D.P. and O.E.A. conceived the idea of the study. D.P. and S.R. supervised analyses. Sv.St. performed QC on the UK Biobank data and wrote the analysis pipeline. K.W. constructed and applied the FUMA pipeline for performing follow-up analyses. J.B. conducted the single cell enrichment analyses. J.H.L and N.S. contributed data. D.P. and I.E.J. wrote the first draft of the paper. All other authors contributed data and critically reviewed the paper.

## Author Information

PF Sullivan reports the following potentially competing financial interests: Lundbeck (advisory committee), Pfizer (Scientific Advisory Board member), and Roche (grant recipient, speaker reimbursement). JHL: Cartana (Scientific Advisor) and Roche (grant recipient). Ole A Andreassen: (Lundbeck) speaker′s honorarium. Stacy Steinberg, Hreinn Stefansson and Kari Stefansson are employees of deCODE Genetics/Amgen. All other authors declare no financial interests or potential conflicts of interest.

Correspondence and requests for materials should be addressed to.

## Online methods

### 1.1 Study Cohorts

#### 1.1.1 PGC-ALZ cohorts

Three non-public datasets (the Norwegian DemGene network, The Swedish Twin Studies of Aging and TwinGene) were meta-analyzed as part of the Alzheimer workgroup initiative of the Psychiatric Genomic Consortium (PGC-ALZ).

We collected genotype data from the Norwegian DemGene Network consisting of 2,224 cases and 1,855 healthy controls. The DemGene Study is a Norwegian network of clinical sites collecting cases from Memory Clinics based on standardised examination of cognitive, functional and behavioural measures and data on progression of most patients. We diagnosed 2,224 cases of AD from 7 studies: the Norwegian Register of persons with Cognitive Symptoms (NorCog), the Progression of Alzheimer′s Disease and Resource use (PADR), the Dementia Study of Western Norway (DemVest), the AHUS study, the Dementia Study in Rural Northern Norway (NordNorge), the HUNT Dementia Stud, the Nursing Home study, and the TrønderBrain study. These cases were diagnosed according to the recommendations from the National Institute on Aging-Alzheimer′s Association (NIA/AA) (AHUS), the NINCDS-ADRDA criteria (DemVest and TrønderBrain) or the ICD-10 research criteria (NorCog, PADR, NordNorge and HUNT). The controls from Norway were obtained through the AHUS, NordNorge, HUNT and TrønderBrain studies. The controls were screened with standardized interview and cognitive tests. Genotypes of the 4079 individuals from the DemGene Study were obtained with Human Omni Express-24 v.1.1 (Illumina Inc., San Diego, CA, USA) at deCODE Genetics (Reykjavik, Iceland). To increase the statistical power of our association analysis, the controls were combined with additional 5786 population controls from Norwegian blood donor samples (Oslo University Hospital, Ullevål Hospital, Oslo) and controls from Thematically Organized Psychosis (TOP) Research Study (between 25-65 years). Control subjects of the TOP Research Study were of Caucasian origin without history of moderate/severe head injury, neurological disorder, mental retardation and were excluded if they or any of their close relatives had a lifetime history of a severe psychiatric disorder, a history of medical problems thought to interfere with brain function or significant illicit drug use.

The Swedish Twin Studies of Aging (STSA) (n cases = 398, n controls = 1079) includes three sub-studies of aging within the Swedish Twin Registry^36^: The Swedish Adoption/Twin Study of Aging (SATSA)^37^, Aging in Women and MEN (GENDER)^38^, and The Study of Dementia in Swedish Twins (HARMONY)^39^. Informed consent was obtained from all participants and the studies were approved by the Regional Ethics Board in Stockholm and the Institutional Review Board at the University of Southern California. DNA was extracted from blood samples and genotyped using Illumina Infinium PsychArray. Alzheimer′s disease patients were diagnosed as part of the studies according to the NINCDS/ADRDA criteria^40^. In addition, information on disease after last study participation was retrieved from three population-based health care registers: The National Patient Register, the Causes of Death Register, and the Prescribed Drug Register.

TwinGene^36^ is a population-based study of older twins drawn from the Swedish Twin Registry. Written informed consent was obtained from all participants and the study was approved by the Regional Ethics Board in Stockholm. DNA was extracted from blood samples and genotyped using Illumina Human OmniExpress for 1791 individuals. Information about Alzheimer′s disease (n cases = 343, n controls = 9070) was extracted from the National Patient Register, the Causes of Death Register, and the Prescribed Drug Register, all of which are population-based health care registers with nationwide coverage.

#### 1.1.2 IGAP

Publically available (http://web.pasteur-lille.fr/en/recherche/u744/igap/igap_download.php) genome-wide association analysis results of the International Genomics of Alzheimer′s Project (IGAP)^4^ were included as one of the four cohorts that were meta-analysed in our effort. IGAP is a large two-stage study based upon genome-wide association studies (GWAS) on individuals of European ancestry. We focused on the results of stage 1, for which IGAP used genotyped and imputed data of 7,055,881 single nucleotide polymorphisms (SNPs) to meta-analyse four previously-published GWAS datasets consisting of 17,008 Alzheimer′s disease cases and 37,154 controls (The European Alzheimer′s disease Initiative EADI, the Alzheimer Disease Genetics Consortium ADGC, the Cohorts for Heart and Aging Research in Genomic Epidemiology consortium CHARGE, the Genetic and Environmental Risk in AD consortium GERAD). As the purpose of stage 2 (11,632 SNPs were genotyped and tested for association in an independent set of 8,572 Alzheimer′s disease cases and 11,312 controls) was replication of the significantly associated loci of stage 1, we limited the inclusion of the summary statistics for our own analyses to stage 1. Written informed consent was obtained from study participants or, for those with substantial cognitive impairment, from a caregiver, legal guardian or other proxy, and the study protocols for all populations were reviewed and approved by the appropriate institutional review boards.

#### 1.1.3 ADSP

The Alzheimer′s Disease Sequencing Project (ADSP) collaboration has the aim to identify novel genetic factors that contribute to AD risk by studying genetic sequencing data. ADSP has made their sequencing data available through the Genotypes and Phenotyps database (dbGaP) under the study accession: phs000572.v7.p (https://www.ncbi.nlm.nih.gov/projects/gap/cgi-bin/study.cgi?study_id=phs000572.v1.p1). We have obtained access to 10,907 individuals (5,771 cases, 5,136 controls) with whole-exome sequencing data to include as the second cohort within our meta-analysis. A substantial proportion of the ADSP individuals were previously also included in IGAP. We applied two strategies to prevent inflated meta-analysis results due to sample overlap: (1) exclusion of ADSP individuals that were duplicates based on genotype data comparison of individual level genetic data between IGAP and ADSP, (2) perform meta-analysis while correcting for cross-study LD score regression intercept (see section 1.4.). To accomplish the first approach we obtained access for all IGAP datasets for which individual level genotype data was available through dbGaP (phs000160.v1.p1 https://www.ncbi.nlm.nih.gov/projects/gap/cgi-bin/study.cgi?study_id=phs000160.v1.p1;phs000219.v1.p1 https://www.ncbi.nlm.nih.gov/projects/gap/cgi-bin/study.cgi?study_id=phs000219.v1.p1;phs000372.v1.p1 https://www.ncbi.nlm.nih.gov/projects/gap/cgi-bin/study.cgi?study_id=phs000372.v1.p1;phs000168.v2.p2 https://www.ncbi.nlm.nih.gov/projects/gap/cgi-bin/study.cgi?study_id=phs000168.v2.p2;phs000234.v1.p1 https://www.ncbi.nlm.nih.gov/projects/gap/cgi-bin/study.cgi?study_id=phs000234.v1.p1)orNIAGADSNG00026 https://www.niagads.org/datasets/ng00026;NG00028 https://www.niagads.org/datasets/ng00028;NG00029 https://www.niagads.org/datasets/ng00029;NG00031 https://www.niagads.org/datasets/ng00030;NG00031 https://www.niagads.org/datasets/ng00031;NG00034 https://www.niagads.org/datasets/ng00034). By calculating identity-by-descent using PLINK^41^, we identified duplicates, which were excluded from the ADSP WES dataset for subsequent analyses.

#### 1.1.1 UK Biobank study

The current study used data from the UK Biobank^42^ (UKB; www.ukbiobank.ac.uk), a large population-based cohort that includes over 500,000 participants and aims to improve insight into a wide variety of health-related determinants and outcomes across the UK. Between 2006 and 2010, approximately 9.2 million invitations to participate in the study were sent to individuals aged 40-69 years who were registered with the National Health Service (NHS) and were living within 25 miles from one of the 22 study research centers. In total, 503,325 participants were recruited in the study, from which we used a subsample of individuals of European ancestry with available phenotypic and genotypic data (*M* age = 56.5, 54.0% female), described in more detail below. Besides phenotypic information obtained from the NHS registries and associated medical records, participants completed an in-person visit at one of the study research centers where extensive self-report data were collected by questionnaire in addition to anthropometric assessments, DNA collection from blood samples, and magnetic resonance imaging of body and brain. All participants provided written informed consent; the UKB received ethical approval from the National Research Ethics Service Committee North West-Haydock (reference 11/NW/0382), and all study procedures were in accordance with the World Medical Association for medical research. Access to the UK Biobank data was obtained under application number 16406.

### 1.2 UKB by proxy phenotype

A proxy phenotype for Alzheimer′s disease case-control status in UKB was assessed as part of the self-report questionnaire administered during the in-person assessment. Participants were asked to report whether their biological mother or father ever suffered from Alzheimer′s disease/dementia, and to report each parent′s current age (or age at death, if applicable). Of 376,113 individuals in our analytic subsample who completed these questions, a diagnosis was reported for 32,327 mothers (8.6%) and 17,014 fathers (4.5%), resulting in 47,793 participants (12.7%) with one or both parents affected. We created a proxy phenotype from these questions to index genetic risk for Alzheimer′s based on parents′ diagnoses. The phenotype was constructed as a linear count of the number of affected biological parents (0, 1, or 2). The contribution for each unaffected parent to this count was weighted by the parent′s age/age at death to account for the fact that they may not yet have passed through the period of risk for this late-onset disease. Specifically, each affected parent contributed one full unit of ″risk″ to the count, while each unaffected parent contributed a proportion of one unit of ″risk″ inversely related to their age. This was calculated as the ratio of parent′s age to age 100 (approximately the 95th percentile for life expectancy in developed countries, such that weight=(100-age)/100. The weight for unaffected parents was capped at 0.32, corresponding to a risk equivalent to that of the maximum population prevalence of AD.^43^ The phenotype thus ranged approximately from 0 to 2, with values near zero when both parents were unaffected (lower for older parents and possible values below zero if both parents were over age 100) and values of two when both parents were affected. Participants who were uncertain or chose not to answer questions about either parent′s disease status or age were excluded from the analyses, resulting in a final N=364,859.

Additional information on Alzheimer′s disease risk was obtained from national medical records linked to participant data. This information pertained to the participants themselves (not their parents), and was extracted from hospital records obtained between 1996 and the present or from national death registries in the case of participants who passed away after initial enrolment in the study, as described in more detail in the UKB resources (http://biobank.ctsu.ox.ac.uk/crystal/refer.cgi?id=146641;; http://biobank.ctsu.ox.ac.uk/crystal/refer.cgi?id=115559). Briefly, primary and secondary diagnoses from inpatient hospital stays and primary and secondary causes of death from death records were recorded using ICD-10 codes. Participants with a diagnosis of ″Alzheimer′s disease″ (diseases of the nervous system chapter; code G30) or “Dementia in Alzheimer′s disease″ (mental and behavioral disorders chapter; code F00) from any record of a hospital stay or as a cause of death were treated as Alzheimer′s cases as given the maximum possible “risk" score of 2, regardless of the affectation status of their parents. The reported rate of Alzheimer′s in parents of cases (27.4%) was more than double that of non-cases (12.7%; ***x***^2^(1)=71.7, *P*=2.45E-17). There were 393 individuals in the analytic subsample classified as affected by these records; due to the small number of cases and the limited representativeness of these types of health records, we used this information to supplement the proxy parent phenotype rather than as a primary outcome. This information reduces the possibility of misclassification in the proxy phenotype method, and also allows us to evaluate the performance of the proxy phenotype method.

### 1.3 Genome-wide association analysis

Except for IGAP (obtained summary statistics), we performed genome-wide association analyses for the ADSP, PGC-ALZ and UKB cohorts. For the UKB dataset, quality control and imputation procedures were slightly different, and therefore described separately in the sections below.

#### 1.3.1a Quality control and imputation procedures for ADSP and PGC-ALZ datasets

Prior to individual quality control steps, all datasets were filtered on a max missingness of 5%. Individuals were excluded when identified as a low quality sample (individual call rate < 0.98), heterozygosity outlier (F +/-.20), gender mismatch (females: F >0.2, males: F < 0.2) when comparing phenotypic and genotypic data, population outlier (defined by principal component boundaries of 1000 Genomes European samples) or being related to another sample (PI_HAT > 0.2). Inclusion criteria for variants encompassed a call rate > 0.98, a case-control missingness difference < 0.02, a Hardy-Weinberg equilibrium p-value < 10×10^−6^ for controls (<10×10^−10^ for cases) and a valid association p-value (excluding the variants with low allele frequencies).

Pre-imputation, the ADSP and PGC-ALZ datasets were checked for palindromic variants with allele frequency close to 0.5, incorrect reference allele definitions, false strand designation and extreme deviations from expected allele frequencies. Subsequently the ADSP and PGC-ALZ datasets were imputed with the 1000 Genomes Phase 3^44^ reference panel. The reported SNPs all have a considerable imputation quality (INFO score>0.591) and variants with a low allele frequency (MAF<0.01) were excluded, resulting in a total of 7508 individuals (4343 cases and 3165 controls) and 260,934 variants for the ADSP cohort and 17477 individuals (2,736 cases and 14,471 controls) and 9,629,492 variants for the PGC-ALZ cohort.

#### 1.3.1b Quality control and imputation for UKB dataset

We used second-release genotype data that were made available by UKB in July 2017. Genotype data collection and processing are described by the UKB in a previous overview paper^45^. DNA was extracted from blood samples and genotyping was completed for 488,366 individuals on one of two Affymetrix genotyping arrays with custom content, the UK BiLEVE Axiom array (N=49,949) or UK Biobank Axiom array (N=438,417), covering 812,428 genetic markers common to both arrays. Of these, 488,377 individuals and 805,426 markers passed the genotype quality control checks conducted by UKB (see http://www.biorxiv.org/content/early/2017/07/20/166298 for details). Samples were excluded for low DNA concentration, call rate < 95%, excess heterozygosity, sex chromosome abnormality, or sample duplication. Variants were excluded if they exhibited poor clustering of allele calls, batch, plate, array, or sex effects, departures from HWE, or discordance between technical replicate samples.

After quality control, the samples were imputed to approximately 92 million SNPs using both the reference panel of the Haplotype Reference Consortium (HRC)^46^ as well as a combined reference panel of the 1000 Genomes Project^44^ and UK10K. As recommended by UKB, we removed variants that were not imputed on the HRC reference panel due to technical errors in the imputation process of the combined panel. We converted imputed variants to hard calls (certainty > 0.9), filtered by imputation quality (INFO score >0.9), and excluded multi-allelic SNPs, indels, SNPs without unique rsID, and SNPs with minor allele frequency (MAF) <0.0001, resulting in 10,847,151 SNPs available for analysis.

For the present study, we selected unrelated individuals of European ancestry. To empirically determine ancestry, we projected genetic principal components from known ancestral populations in the 1000 Genomes Project onto the UKB genotypes and assigned individuals to the continental ancestral superpopulation with the closest Mahalanobis distance.^47^ Within-ancestry principal components were created using FlashPCA2^48^ to correct for any residual population stratification within the European ancestry subset. Unrelated individuals (less than 3rd degree relatives, as indicated by genomic relatedness coefficients calculated by UKB) were selected by sequentially removing participants with the greatest number of relatives until no related pairs remained. After applying these filtering criteria and removing any participants with missing phenotypic or covariate data and participants who withdrew consent, 364,859 individuals remained for analysis in the UKB sample.

##### 1.3.2 Single-marker association analysis

Genome-wide association analysis (GWAS) for the ADSP, PGC-ALZ and UKB datasets was performed in PLINK^41^, using logistic regression for dichotomous phenotypes (cases versus controls for ADSP and PGC-ALZ cohorts), and linear regression for phenotypes analysed as continuous outcomes (by proxy parental AD phenotype for UKB cohort). For the ADSP and PGC ALZ cohorts, association tests were adjusted for gender, batch (if applicable), and the first 4 principal components. Twenty principal components were calculated, and depending on the dataset being tested, additional principal components (on top of the standard inclusion of 4 PCAs) were added if significantly associated to the phenotype. Furthermore, for the PGC-ALZ cohorts age was included as a covariate. For 4,537 controls of the DemGene cohort, no detailed age information was available, besides the age range the subjects were in (20-45 years). We therefore set the age of these individuals conservatively to 20 years. For the ADSP dataset, age was not included as a covariate due to the enrichment for older controls (mean age cases = 73.1 years (SE=7.8); mean age controls = 86.1 years (SE=4.5)) in their collection procedures. Correcting for age in ADSP would remove a substantial part of genuine association signals (e.g. well-established *APOE* locus rs11556505 is strongly associated to AD (*P*=1.08×10^−99^), which is lost when correcting for age (*P*=0.0054). For the UKB dataset, 12 components were included as covariates, as well as age, sex, genotyping array, and assessment centre. We used the genome wide threshold for significance of P<5×10-8).

##### 1.3.3 Multivariate genome-wide meta-analysis

Two meta-analyses were performed, including: 1) cohorts with case-control phenotypes (IGAP, ADSP and PGC-ALZ datasets), 2) all cohorts, also including the by proxy phenotype of UKB.

The per SNP test statistics is defined by

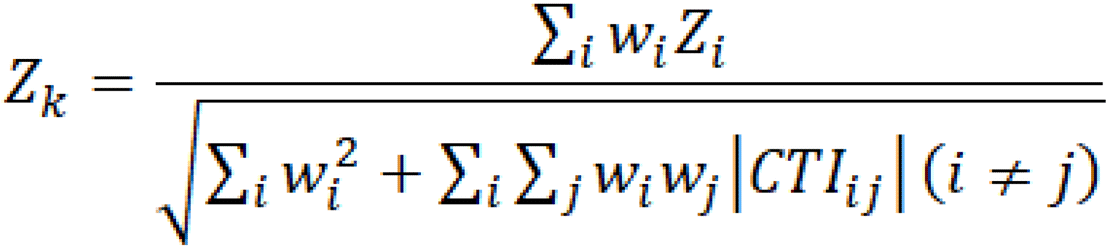

where *w_i_* and *Z_i_* are the squared root of the sample size and the test statistics of SNP *k* in cohort *i*, respectively. CTI is the cross trait LD score intercept estimated by LDSC using genome-wide summary statistics as

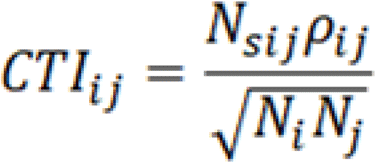

where *N_sij_* and *r_ij_* are the number of overlapping samples and the phenotypic correlation between cohort *i* and *j,* respectively.^14^ The test statistics per SNP per GWAS were converted from the P-value by using the sign of either beta or odds ratio. When direction is aligned the conversion is two-sided. To avoid infinite values, we replaced P-value 1 with 0.999999 and P value < 1e-323 to 1e-323 (the minimum >0 value in Python).

The effective sample size (*N_eff_*) is computed for each SNP *k* from the matrix *M,* containing the sample size *N_i_* of each cohort *i* on the diagonal and the estimated number of shared data points 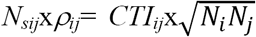 for each pair of cohorts *i* and *j* as the off-diagonal values. *N_eff_* is computed recursively as follows. Starting with the first cohort in *M*, *N_eff_* is first increased by *M_1,1_,* corresponding to the sample size of that cohort. The proportion of samples shared between cohort 1 and each other cohort *j* is then computed as *p_1,j_* = *M_1,j_/M_j,j_,* and *M* is then adjusted to remove this overlap, multiplying all values in each column *j* by *1-p_i,j_.* This amounts to reducing the sample size of each other cohort *j* by the number of samples it shares with cohort 1, and reducing the shared samples between cohort *j* and subsequent cohorts by the same proportion. After this, the first row and column of *M* are discarded, and the same process is applied to the new *M* matrix. This is repeated until *M* is empty. The script for the multivariate GWAS is available from https://github.com/Kyoko-wtnb/mvGWAMA.

#### 1.5 Replication of meta-analysis result in an Icelandic sample

The study group included 6,593 Alzheimer′s disease cases (4,923 of whom were chip-typed) and 174,289 controls (88,581 of whom were chip-typed). In 16% of patients, the diagnosis of Alzheimer′s disease was established at the Memory Clinic of the University Hospital according to the criteria for definite, probable, or possible Alzheimer′s disease of the National Institute of Neurological and Communicative Disorders and Stroke and the Alzheimer′s Disease and Related Disorders Association (NINCDS-ADRDA). In 77% of patients, the diagnosis has been registered according to the criteria for code 331.0 in ICD-9, or for F00 and G30 in ICD-10 in health records. Seven percent of the patients were identified in the Directorate of Health medication database as having been prescribed Donepezil (Aricept). The controls were drawn from various research projects at deCODE Genetics.

The study was approved by the National Bioethics Committee and the Icelandic Data Protection Authority. Written informed consent was obtained from all participants or their guardians before blood samples were drawn. All sample identifiers were encrypted in accordance with the regulations of the Icelandic Data Protection Authority.

Chip-typing and long-range phasing of 155,250 individuals was carried out as described previously.^21^ Imputation of the variants found in 28,075 whole-genome sequenced individuals into the chip-typed individuals and 285,664 close relatives was performed as detailed earlier.^21^ Association analysis was carried out using logistic regression with Alzheimer′s disease status as the response and genotype counts and a set of nuisance variables including sex, county of birth, and current age as predictors.^22^ Correction for inflation of test statistics due to relatedness and population stratification was performed using the intercept estimate from LD score regression^14^ (1.29).

#### 1.6 Genomic risk loci definition

We used FUMA^26^, an online platform for functional mapping and annotation of genetic variants, to define genomic risk loci and obtain functional information of relevant SNPs in these loci. We first identified independent significant SNPs that have a genome-wide significant P-value (<5×10^−8^) and are independent from each other at *r^2^* <0.6. These SNPs were further represented by lead SNPs, which are a subset of the independent significant SNPs that are in approximate linkage equilibrium with each other at *r^2^*>0.6. We then defined associated genomic risk loci by merging any physically overlapping lead SNPs (LD blocks <250kb apart). Borders of the genomic risk loci were defined by identifying all SNPs in LD (*r^2^*>0.6) with one of the independent significant SNPs in the locus, and the region containing all these candidate SNPs was considered to be a single independent genomic risk locus. LD information was calculated using the UK Biobank genotype data as a reference.

#### 1.7 Cohort Heritability and Genetic Correlation

LD score regression^14^ was used to estimate genomic inflation and heritability of the AD in each of the 7 cohorts (PGC-ALZ, ADSP, IGAP, UKB, DemGene, STSA, TwinGene) using their post-quality control summary statistics, and to estimate the cross-cohort genetic correlations.^49^ Pre-calculated LD scores from the 1000 Genomes European reference population were obtained from https://data.broadinstitute.org/alkesgroup/LDSCORE/. Genetic correlations were calculated on HapMap3 SNPs only. LD score regression was also used on the case-control and by-proxy phenotype result to estimate heritability and genetic correlations for the two phenotype definitions.

#### 1.8 Polygenic risk scoring

We calculated polygenic scores (PGS) based on the SNP effect sizes estimated in the meta-analyses. PGS were calculated using an independent genotype dataset of 761 individuals (379 cases and 382 controls) from the ADDNeuroMed study.^50^ The same QC and imputation approach was applied as for the other datasets with genotype-level data (see Method section 1.3.1a). PRSice PGS were calculated on hard-called imputed genotypes using *P*-value thresholds from 0.0 to 0.5 in steps ranging from 5×10^−8^ to 0.001. The explained variance (Δ*R^2^*) was derived from a linear model in which the AD phenotype was regressed on each PGS while controlling for the same covariates as in each cohort-specific GWAS, compared to a linear model with GWAS covariates only.

#### 1.9 Stratified Heritability

To test whether specific categories of SNP annotations were enriched for heritability, we partitioned the SNP heritability for binary annotations using stratified LD score regression (https://github.com/bulik/ldsc)^14^. Heritability enrichment was calculated as the proportion of heritability explained by a SNP category divided by the proportion of SNPs that are in that category. Partitioned heritability was computed by 28 functional annotation categories, by minor allele frequency (MAF) in six percentile bins and by 22 chromosomes. Annotations for binary categories of functional genomic characteristics (e.g. coding or regulatory regions) were obtained from the LD score website (https://github.com/bulik/ldsc). The Bonferroni-corrected significance threshold for 56 annotations was set at: *P*<0.05/56=8.93×10^−4^.

#### 1.10 Functional Annotation of SNPs

Functional annotation of SNPs implicated in the meta-analysis was performed using FUMA^26^ (http://fuma.ctglab.nl/). We selected all *candidate SNPs* in the associated genomic loci having an *r^2^*≧0.6 with one of the independent significant SNPs (see above), a *P*-value (*P*<1×10^−8^) and a MAF>0.0001 for annotations. Functional consequences for these SNPs were obtained by matching SNPs′ chromosome, base-pair position, and reference and alternative alleles to databases containing known functional annotations, including ANNOVAR^51^ categories, Combined Annotation Dependent Depletion (CADD) scores^24^, RegulomeDB^52^ (RDB) scores, and chromatin states^53,54^. ANNOVAR annotates the functional consequence of SNPs on genes (e.g. intron, exon, intergenic). CADD scores predict how deleterious the effect of a SNP with higher scores referring to higher deleteriousness. A CADD score above 12.37 is the threshold to be potentially pathogenic^55^. The RegulomeDB score is a categorical score based on information from expression quantitative trait loci (eQTLs) and chromatin marks, ranging from 1a to 7 with lower scores indicating an increased likelihood of having a regulatory function. Scores are as follows: 1a=eQTL + Transciption Factor (TF) binding + matched TF motif + matched DNase Footprint + DNase peak; 1b=eQTL + TF binding + any motif + DNase Footprint + DNase peak; 1c=eQTL + TF binding + matched TF motif + DNase peak; 1d=eQTL + TF binding + any motif + DNase peak; 1e=eQTL + TF binding + matched TF motif; 1f=eQTL + TF binding / DNase peak; 2a=TF binding + matched TF motif + matched DNase Footprint + DNase peak; 2b=TF binding + any motif + DNase Footprint + DNase peak; 2c=TF binding + matched TF motif + DNase peak; 3a=TF binding + any motif + DNase peak; 3b=TF binding + matched TF motif; 4=TF binding + DNase peak; 5=TF binding or DNase peak; 6=other;7=None. The chromatin state represents the accessibility of genomic regions (every 200bp) with 15 categorical states predicted by a hidden Markov model based on 5 chromatin marks for 127 epigenomes in the Roadmap Epigenomics Project^39^. A lower state indicates higher accessibility, with states 1-7 referring to open chromatin states. We annotated the minimum chromatin state across tissues to SNPs. The 15 core chromatin states as suggested by Roadmap are as follows: 1=Active Transcription Start Site (TSS); 2=Flanking Active TSS; 3=Transcription at gene 5′ and 3′; 4=Strong transcription; 5= Weak Transcription; 6=Genic enhancers; 7=Enhancers; 8=Zinc finger genes & repeats; 9=Heterochromatic; 10=Bivalent/Poised TSS; 11=Flanking Bivalent/Poised TSS/Enh; 12=Bivalent Enhancer; 13=Repressed PolyComb; 14=Weak Repressed PolyComb; 15=Quiescent/Low.

#### 1.11 Gene-mapping

Genome-wide significant loci obtained by GWAS were mapped to genes in FUMA^26^ using three strategies:

1. Positional mapping maps SNPs to genes based on physical distance (within a 10kb window) from known protein coding genes in the human reference assembly (GRCh37/hg19).
2. eQTL mapping maps SNPs to genes with which they show a significant eQTL association (i.e. allelic variation at the SNP is associated with the expression level of that gene). eQTL mapping uses information from 45 tissue types in 3 data repositories (GTEx^56^, Blood eQTL browser^57^, BIOS QTL browser^58^), and is based on cis-eQTLs which can map SNPs to genes up to 1Mb apart. We used a false discovery rate (FDR) of 0.05 to define significant eQTL associations.
3. Chromatin interaction mapping was performed to map SNPs to genes when there is a three-dimensional DNA-DNA interaction between the SNP region and another gene region. Chromatin interaction mapping can involve long-range interactions as it does not have a distance boundary. FUMA currently contains Hi-C data of 14 tissue types from the study of Schmitt et al^59^. Since chromatin interactions are often defined in a certain resolution, such as 40kb, an interacting region can span multiple genes. If a SNPs is located in a region that interacts with a region containing multiple genes, it will be mapped to each of those genes. To further prioritize candidate genes, we selected only genes mapped by chromatin interaction in which one region involved in the interaction overlaps with a predicted enhancer region in any of the 111 tissue/cell types from the Roadmap Epigenomics Project^54^ and the other region is located in a gene promoter region (250bp up and 500bp downstream of the transcription start site and also predicted by Roadmap to be a promoter region). This method reduces the number of genes mapped but increases the likelihood that those identified will have a plausible biological function. We used a FDR of 1×10^−5^ to define significant interactions, based on previous recommendations^44^ modified to account for the differences in cell lines used here.

#### 1.12 Gene-based analysis

To account for the distinct types of genetic data in this study, genotype array (PGC-ALZ, IGAP, UKB) and whole-exome sequencing data (ADSP), we first performed two gene-based genome wide association analysis (GWGAS) using MAGMA^30^, followed by a meta-analysis. SNP-based P values from the meta-analysis of the 3 genotype-array-based datasets were used as input for the first GWGAS, while the unimputed individual-level sequence data of ADSP was used as input for the second GWGAS. 18,233 protein-coding genes (each containing at least one SNP in the GWAS) from the NCBI 37.3 gene definitions were used as basis for GWGAS in MAGMA. Bonferroni correction was applied to correct for multiple testing (P<2.74×10^−6^).

#### 1.13 Gene-set analysis

Results from the GWGAS analyses were used to test for association in 7,087 predefined gene sets of four types:

1. 6,994 curated gene-sets representing known biological and metabolic pathways derived from Gene Ontology (5917 gene-sets), Biocarta (217 gene-sets), KEGG (186 gene-sets), Reactome (674 gene-sets) catalogued by and obtained from the MsigDB version 6.1^60^ (http://software.broadinstitute.org/gsea/msigdb/collections.jsp)
2. Gene expression values from 54 (53 + 1 calculated 1^st^ PC of three tissue subtypes) tissues obtained from GTEx^56^, log2 transformed with pseudocount 1 after winsorization at 50 and averaged per tissue.
3. Cell-type specific expression in 173 types of brain cells (24 broad categories of cell types, ′level 1′ and 129 specific categories of cell types ′level 2′), which were calculated following the method described in ^32^. Briefly, brain cell-type expression data was drawn from single-cell RNA sequencing data from mouse brains. For each gene, the value for each cell-type was calculated by dividing the mean Unique Molecular Identifier (UMI) counts for the given cell type by the summed mean UMI counts across all cell types. Single-cell gene-sets were derived by grouping genes into 40 equal bins based on specificity of expression.
4. Nucleus specific gene expression of 15 distinct human brain cell-types from the study described in^61^. The value for each cell-type was calculated with the same method as explained in point 3 above.

These gene-sets were tested using MAGMA. We computed competitive P-values, which represent the test of association for a specific gene-set compared with genes not in the gene set to correct for baseline level of genetic association in the data. The Bonferroni-corrected significance threshold was 0.05/7,087 gene-sets=7.06×10^−6^. The suggestive significance threshold was defined by the number of tests within the category. Conditional analyses were performed as a follow-up using MAGMA to test whether each significant association observed was independent of all others and of *APOE* (a gene-set including all genes within genomic region chr19:45,020,859-45,844,508). Furthermore, the association between each of the significant gene-sets was tested conditional on each of the other significantly associated gene sets. Gene-sets that retained their association after correcting for other sets were considered to represent independent signals. We note that this is not a test of association per se, but rather a strategy to identify, among gene-sets with known significant associations and overlap in genes, which set (s) are responsible for driving the observed association.

#### 1.14 Cross-Trait Genetic Correlation

Genetic correlations (*r_g_*) between AD and 41 phenotypes were computed using LD score regression^14^, as described above, based on GWAS summary statistics obtained from publicly available databases (http://www.med.unc.edu/pgc/results-and-downloads; http://ldsc.broadinstitute.org/; **Supplementary Table 19**). The Bonferroni-corrected significance threshold was 0.05/41 traits=1.22×10^−3^.

#### 1.15 Mendelian Randomisation

To infer credible causal associations between AD and traits that are genetically correlated with AD, we performed Generalised Summary-data based Mendelian Randomisation^31^ (GSMR; http://cnsgenomics.com/software/gsmr/). This method utilizes summary-level data to test for putative causal associations between a risk factor (exposure) and an outcome by using independent genome-wide significant SNPs as instrumental variables as an index of the exposure. HEIDI-outlier detection was used to filter genetic instruments that showed clear pleiotropic effects on the exposure phenotype and the outcome phenotype. We used a threshold p-value of 0.01 for the outlier detection analysis in HEIDI, which removes 1% of SNPs by chance if there is no pleiotropic effect. To test for a potential causal effect of various outcomes on risk for AD, we selected phenotypes in non-overlapping samples that showed (suggestive) significant (*P*<0.05) genetic correlations (*r_g_*) with AD. With this method it is typical to test for bi-directional causation by repeating the analyses while switching the role of the exposure and the outcome; however, because AD is a late-onset disease, it makes little sense to estimate its causal effect on outcomes that develop earlier in life, particularly when the summary statistics for these outcomes were derived mostly from younger samples than those of AD cases. Therefore, we conducted these analyses only in one direction. For genetically correlated phenotypes, we selected independent (*r^2^*=<0.1), GWS lead SNPs as instrumental variables in the analyses. The method estimates a putative causal effect of the exposure on the outcome (*b_xy_*) as a function of the relationship between the SNPs′ effects on the exposure (b_zx_) and the SNPs′ effects on the outcome (*b_zy_*), given the assumption that the effect of non pleiotropic SNPs on an exposure (x) should be related to their effect on the outcome (y) in an independent sample only via mediation through the phenotypic causal pathway (*b_xy_*). The estimated causal effect coefficients (*b_xy_*) are approximately equal to the natural log odds ratio (OR)^31^ for a case-control trait. An OR of 2 can be interpreted as a doubled risk compared to the population prevalence of a binary trait for every SD increase in the exposure trait. For quantitative traits the *b_zx_* and *b_zy_* can be interpreted as a one standard deviation increase explained in the outcome trait for every SD increase in the exposure trait. This method can help differentiate the causal direction of association between two traits, but cannot make any statement about the intermediate mechanisms involved in any potential causal process.

## Data availability

Summary statistics will be made available for download upon publication (https://ctg.cncr.nl).

---

**Supplementary Figure 1.**
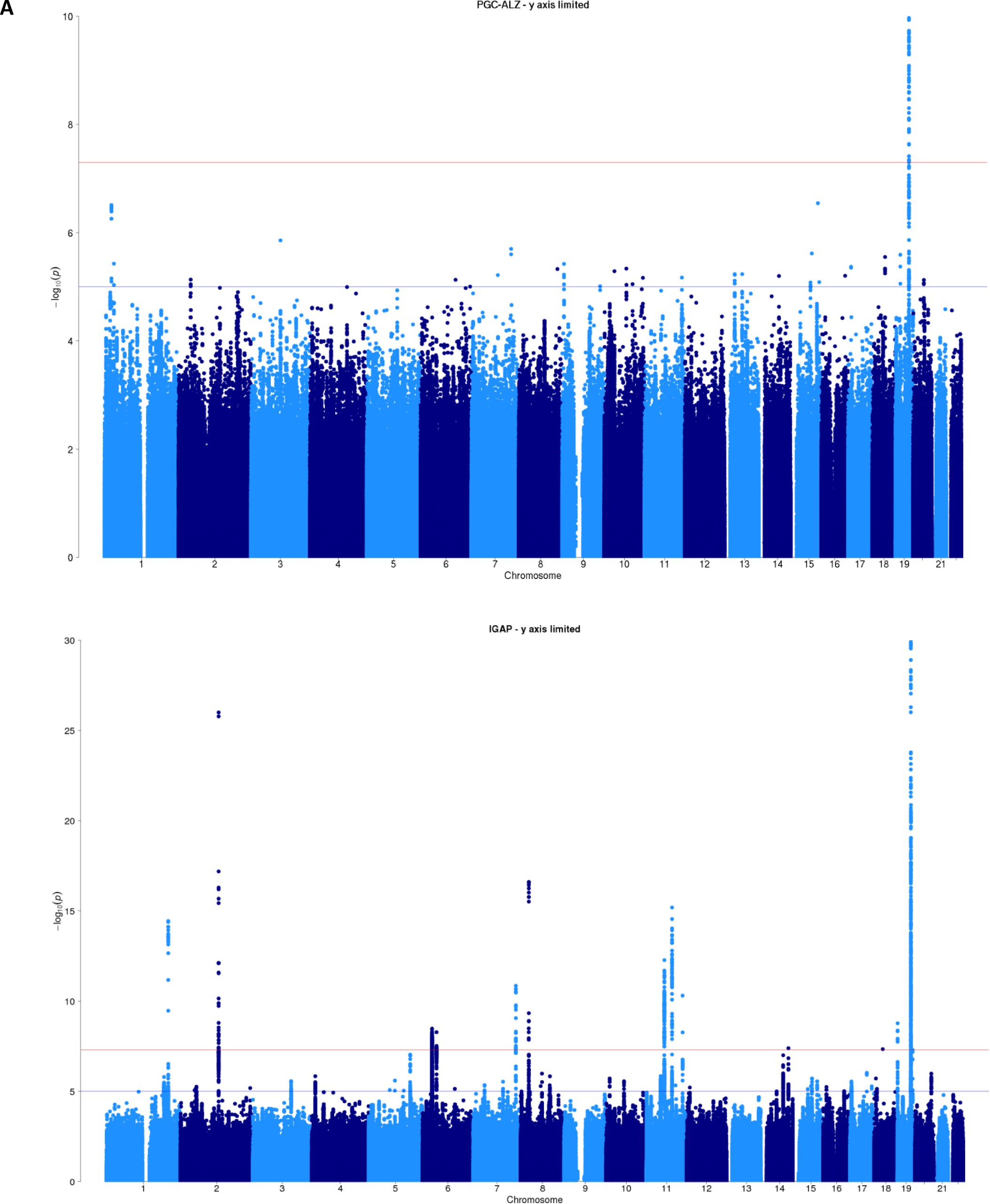
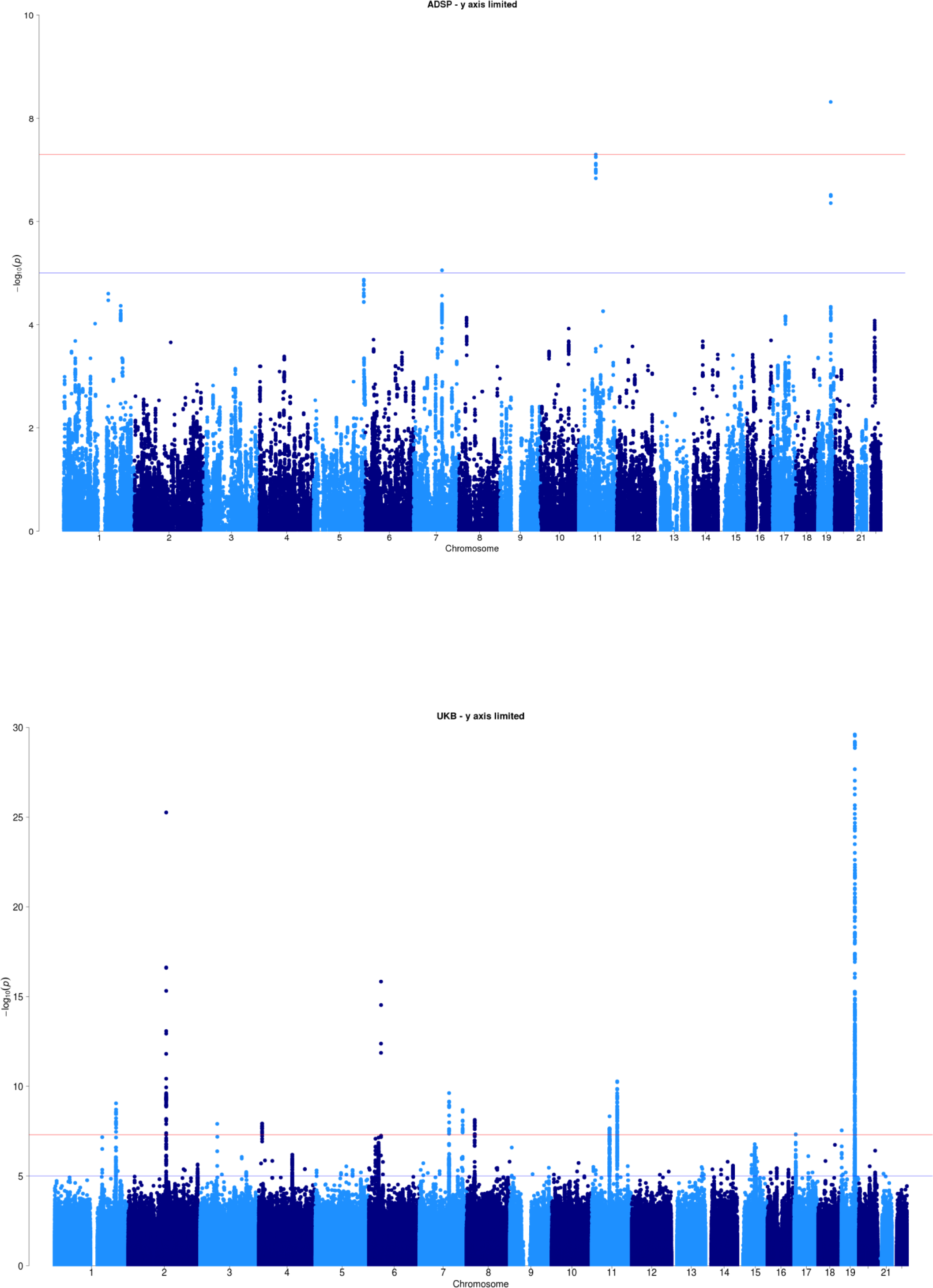
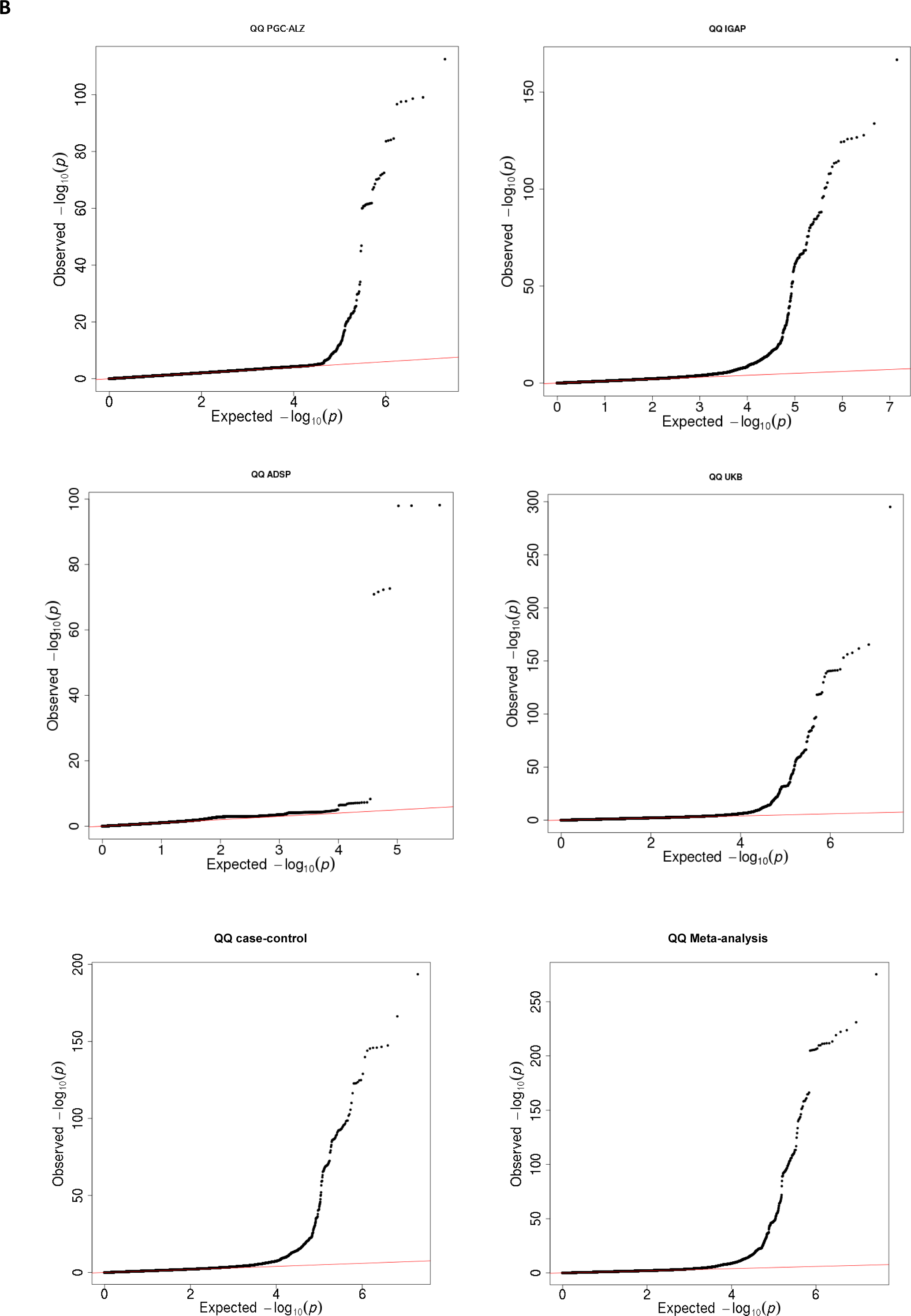
Manhattan and QQ plots of single variant association results per main cohort. For each cohort, Manhattan and QQ plots are shown. A) The Manhattan plot displays all associations per variant ordered according to their genomic position on the x-axis and showing the strength of the association with the −log10 transformed P-values on the y-axis. The y-axis is limited to enable visualization of *non-APOE* loci. B) The QQ plot displays the expected −log10 transformed *p-* values on the x-axis and the observed −log10 transformed p-values on the y-axis.

**Supplementary Figure 2.**
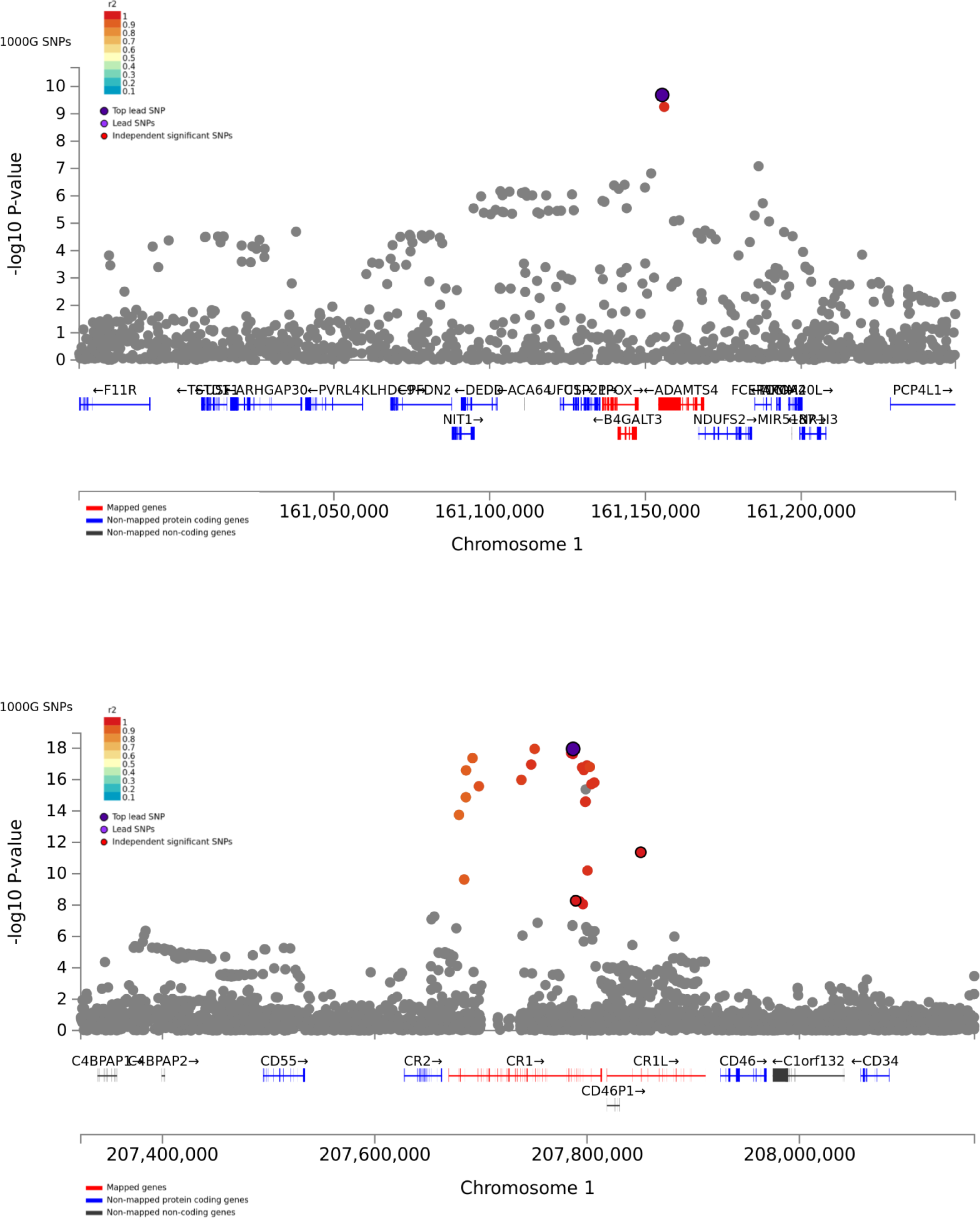
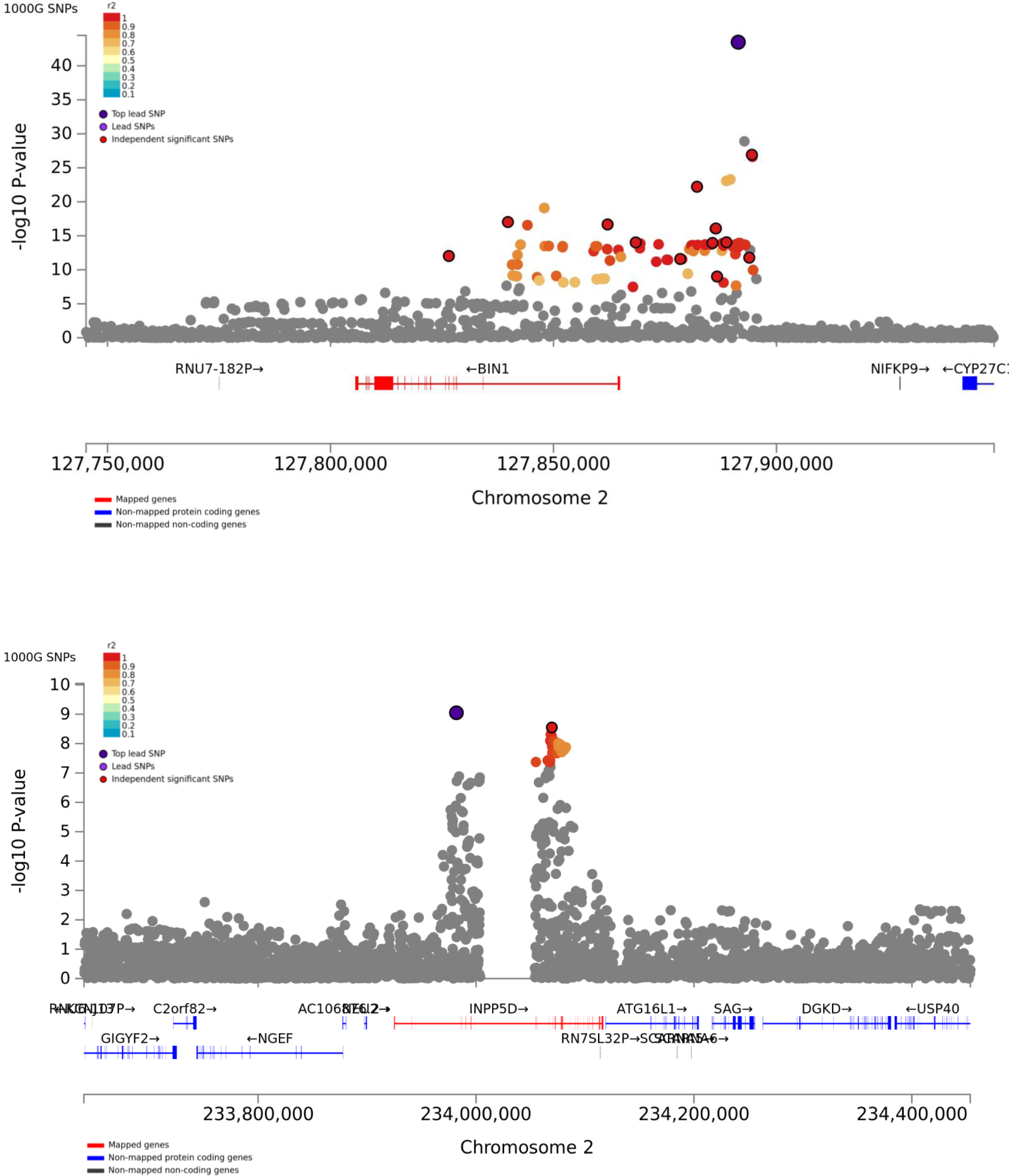
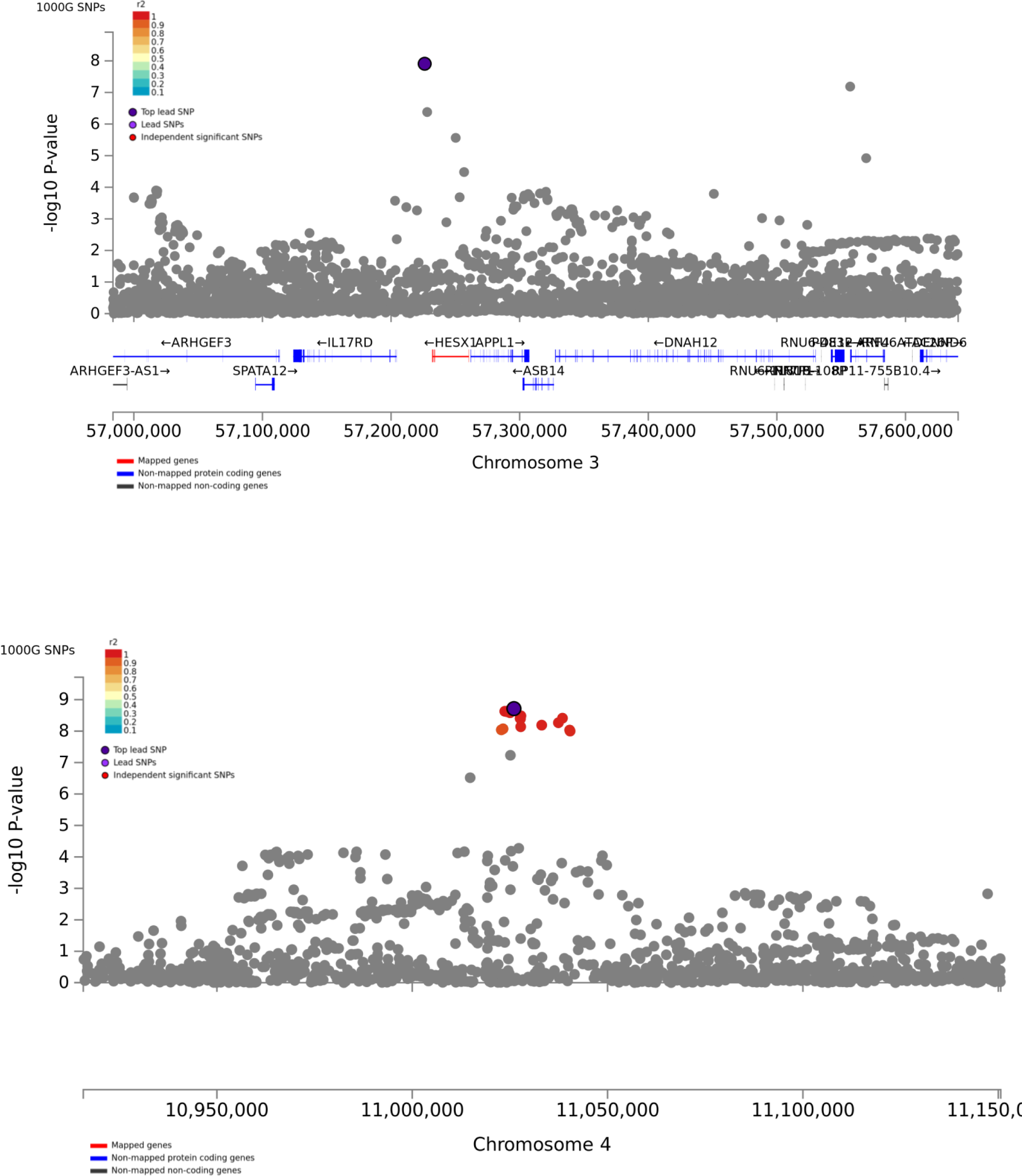
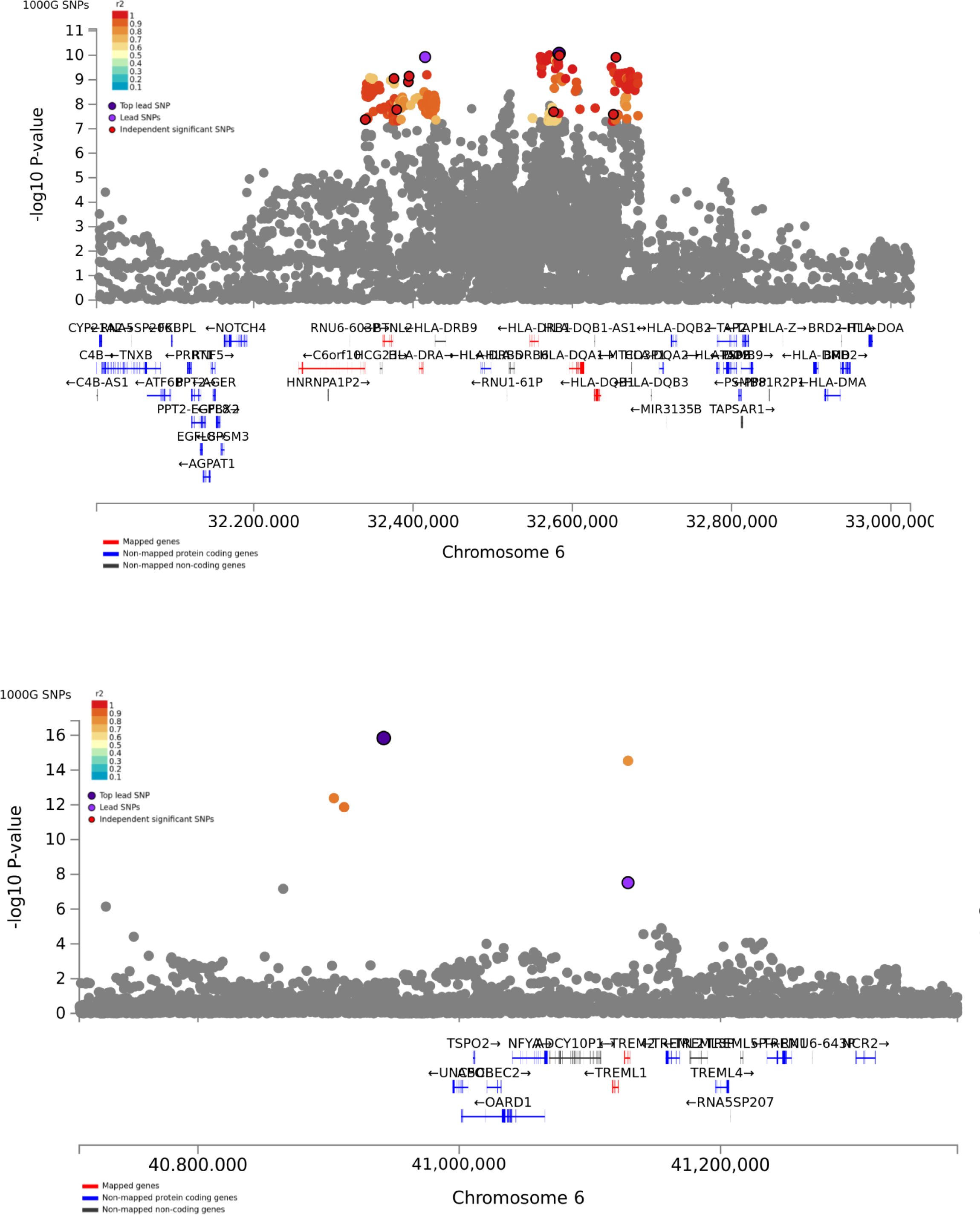
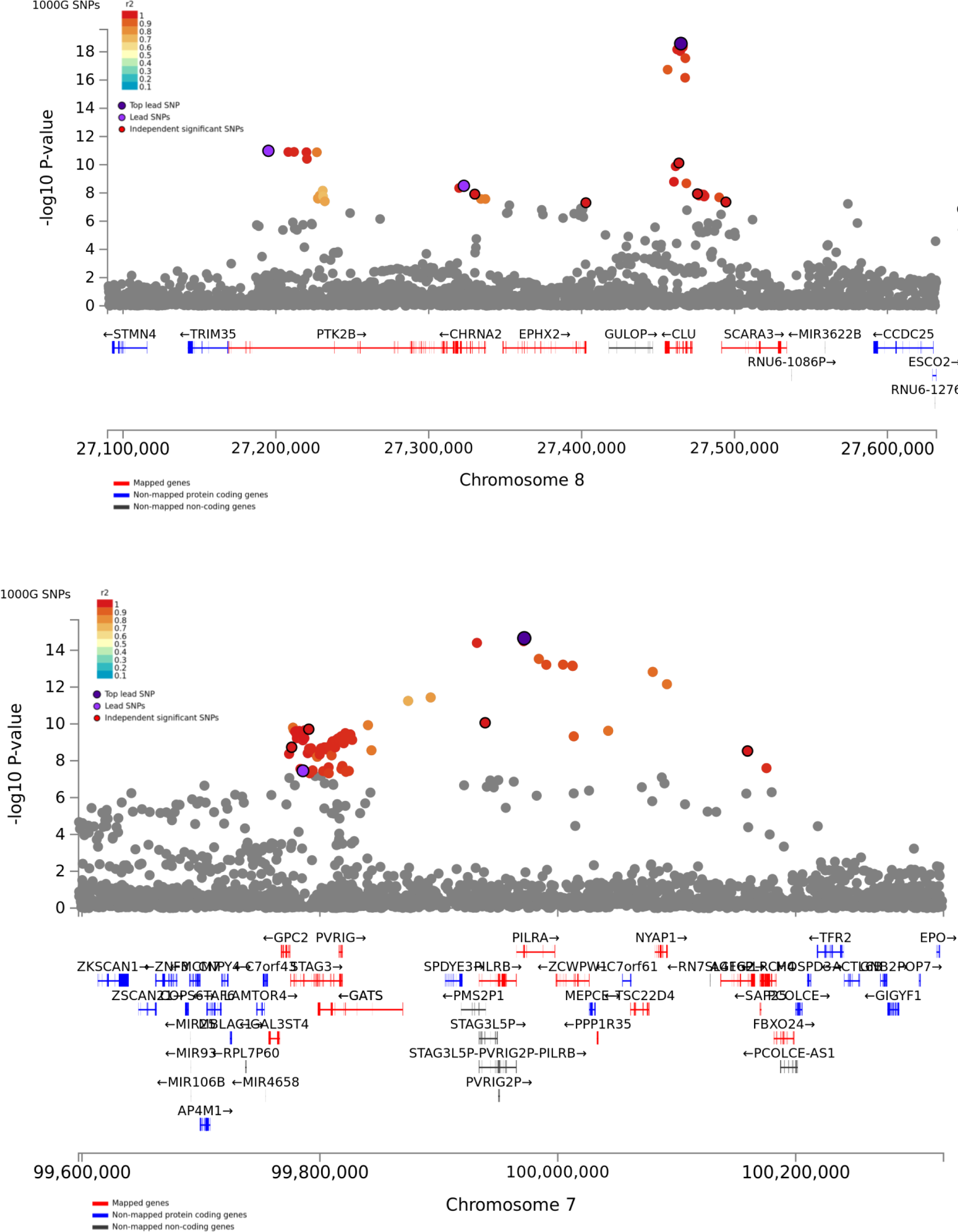
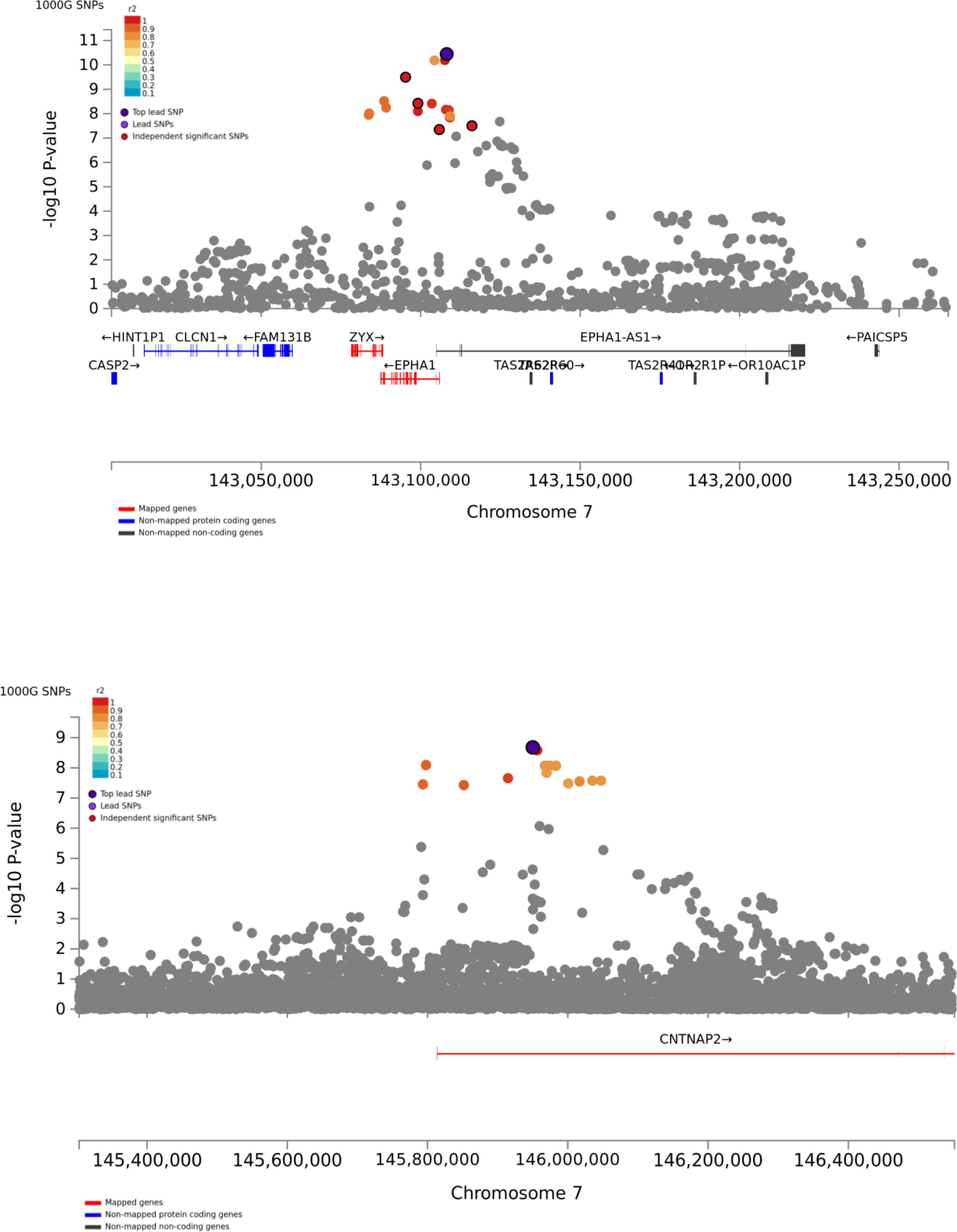
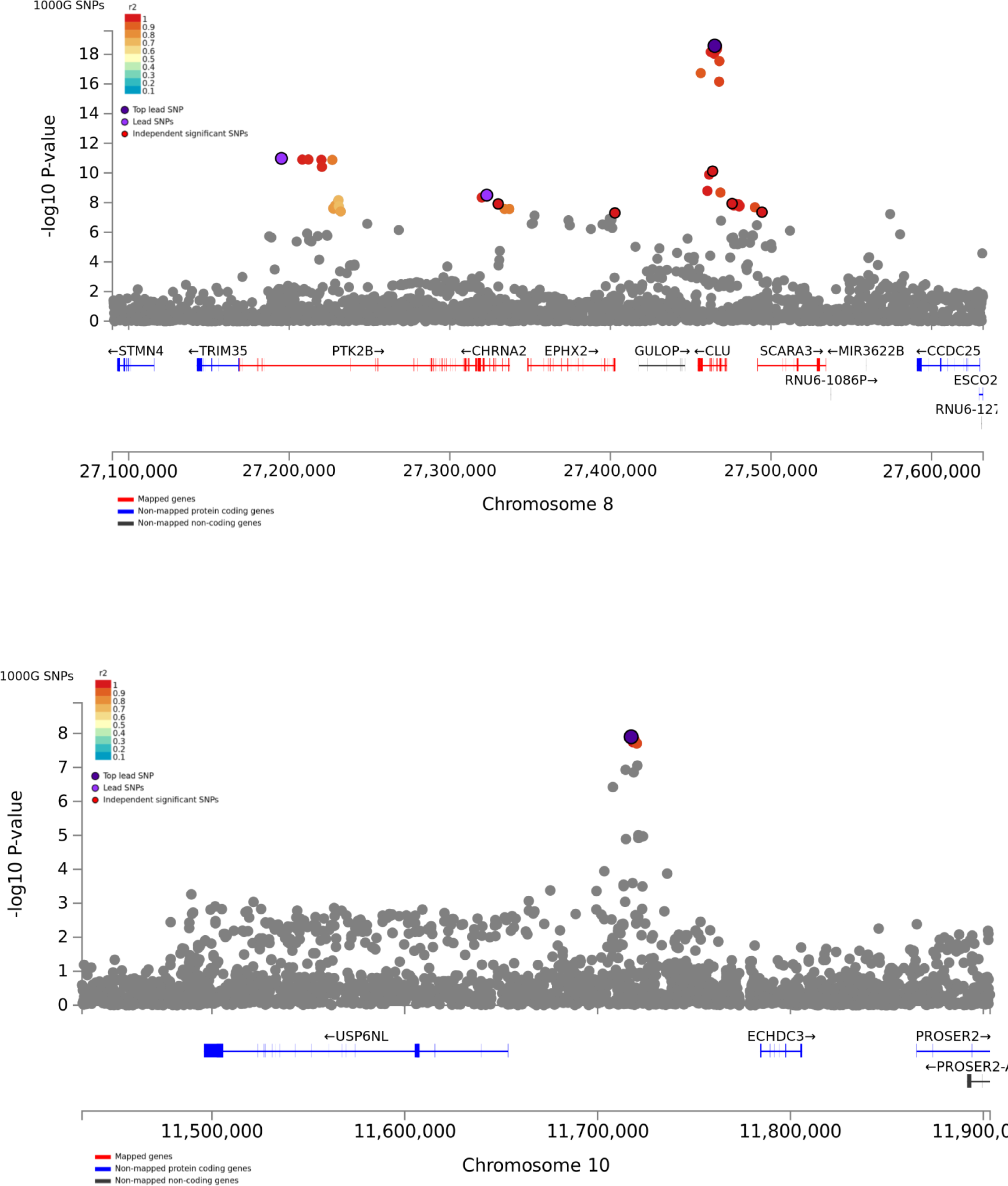
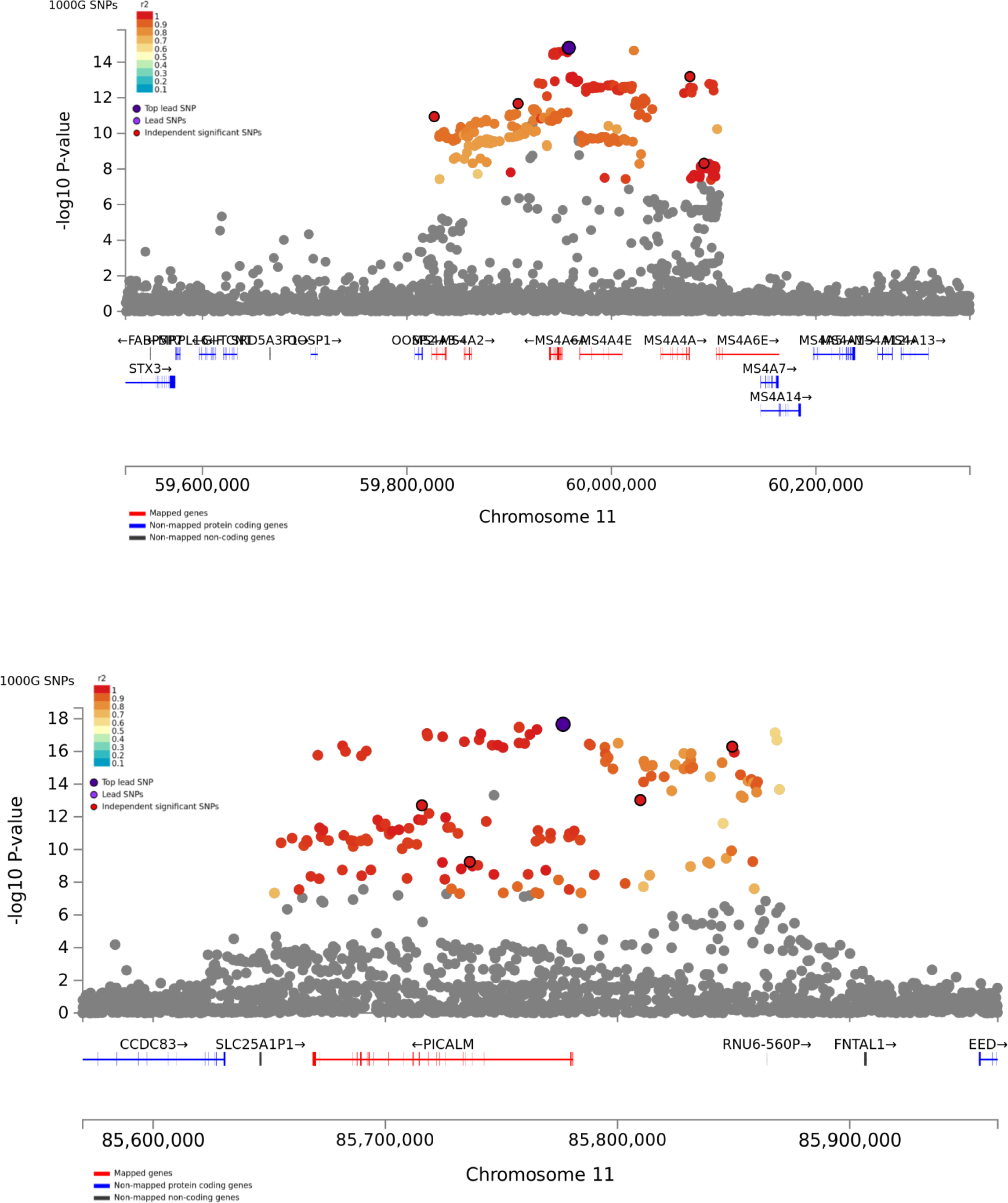
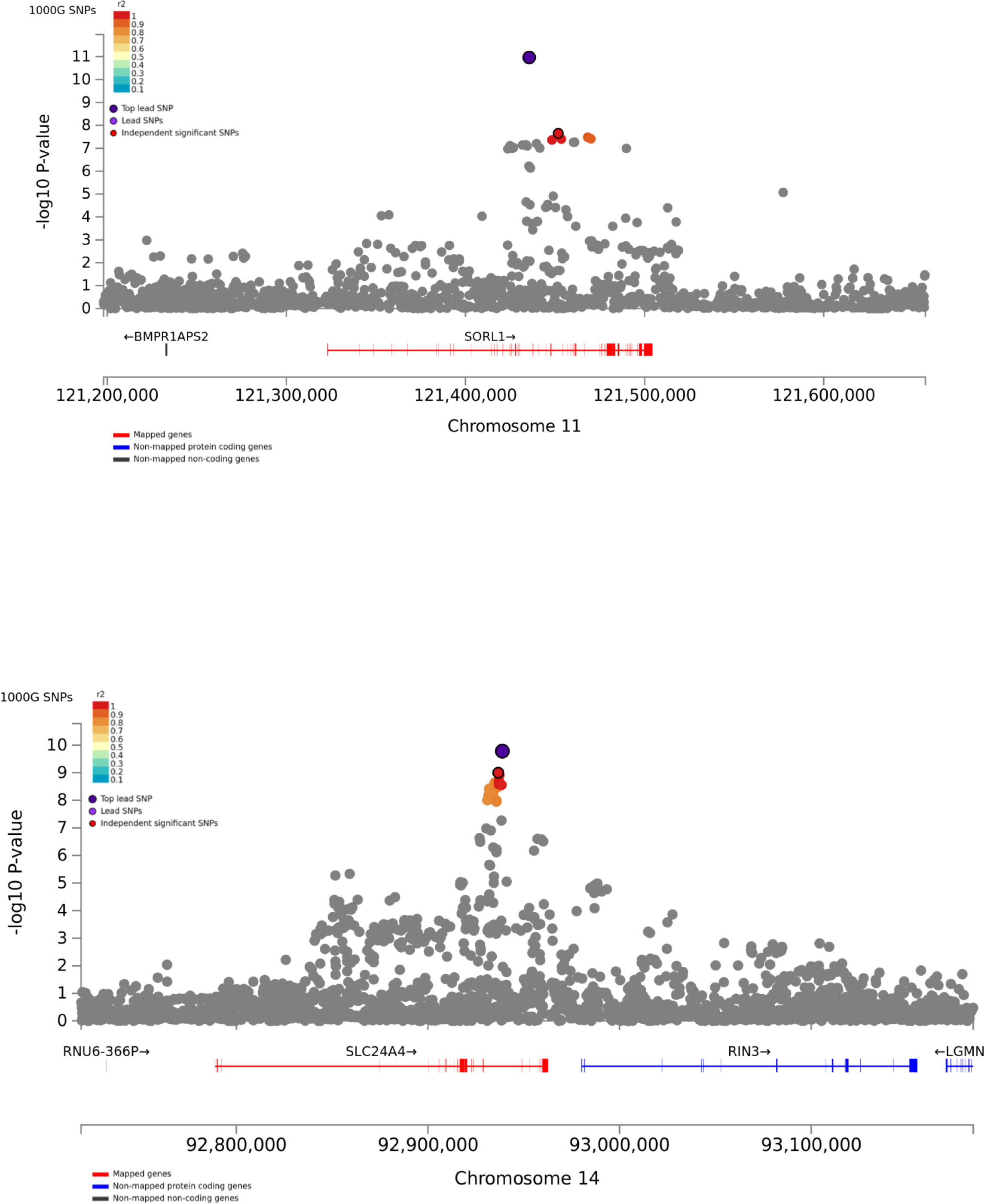
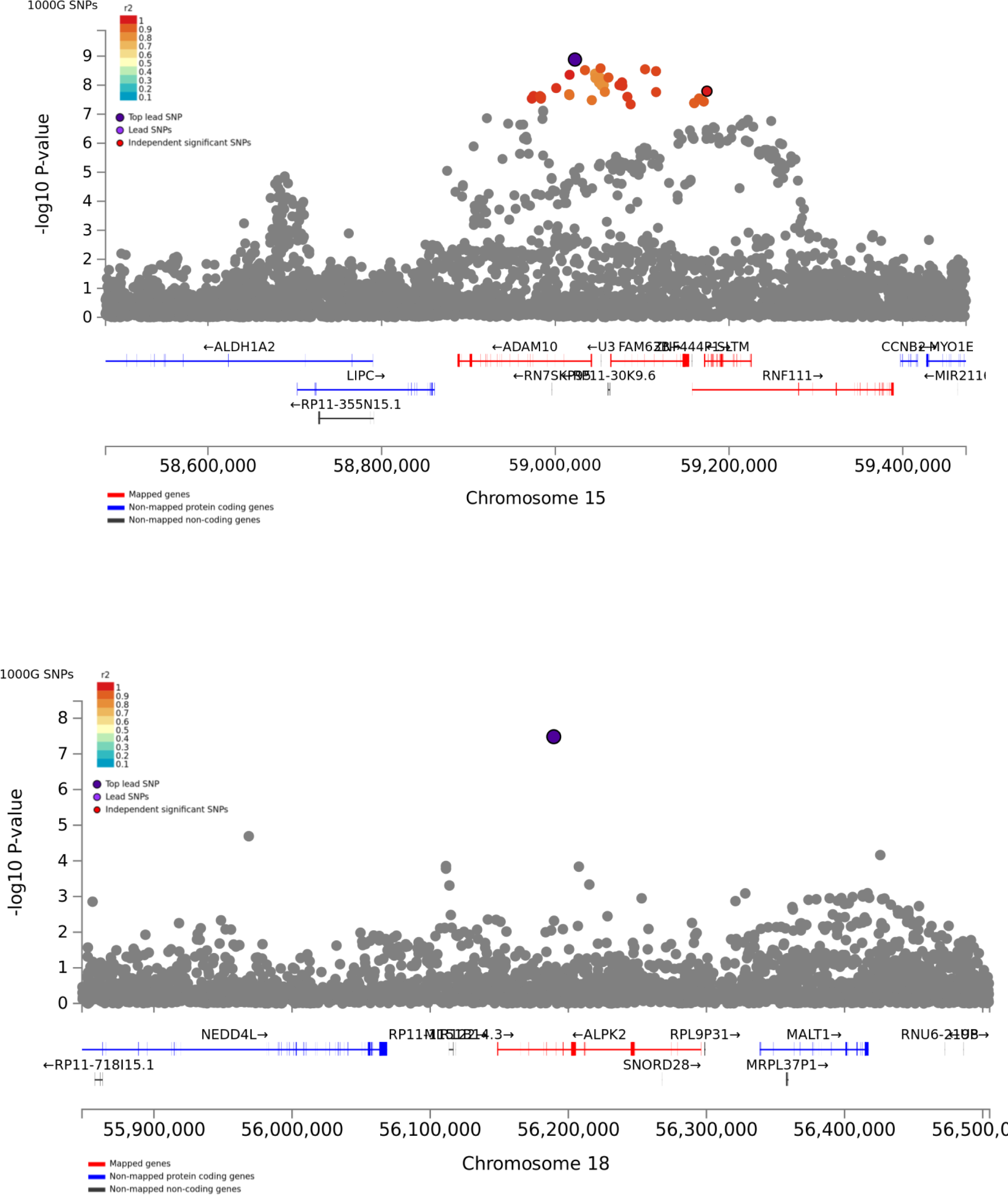
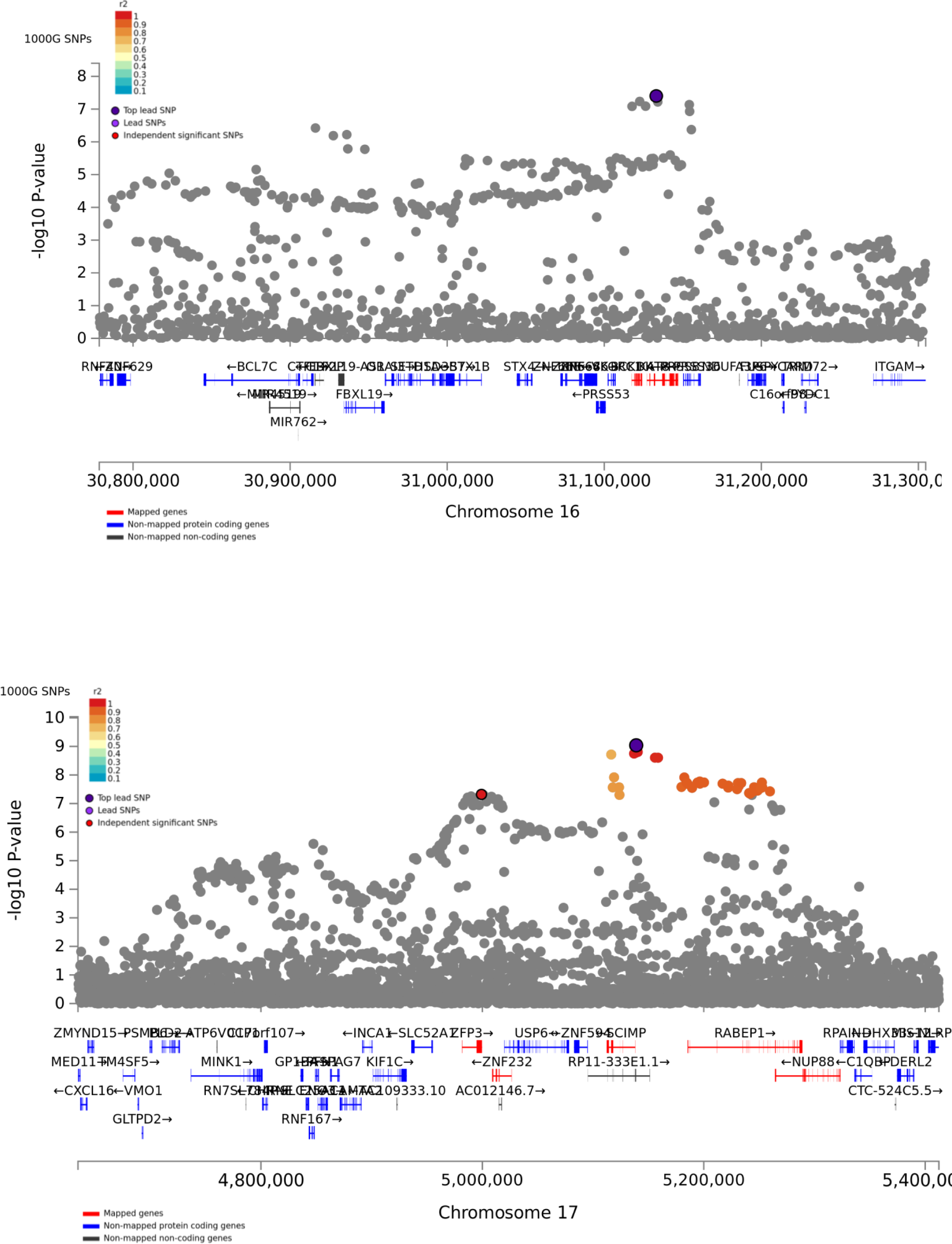
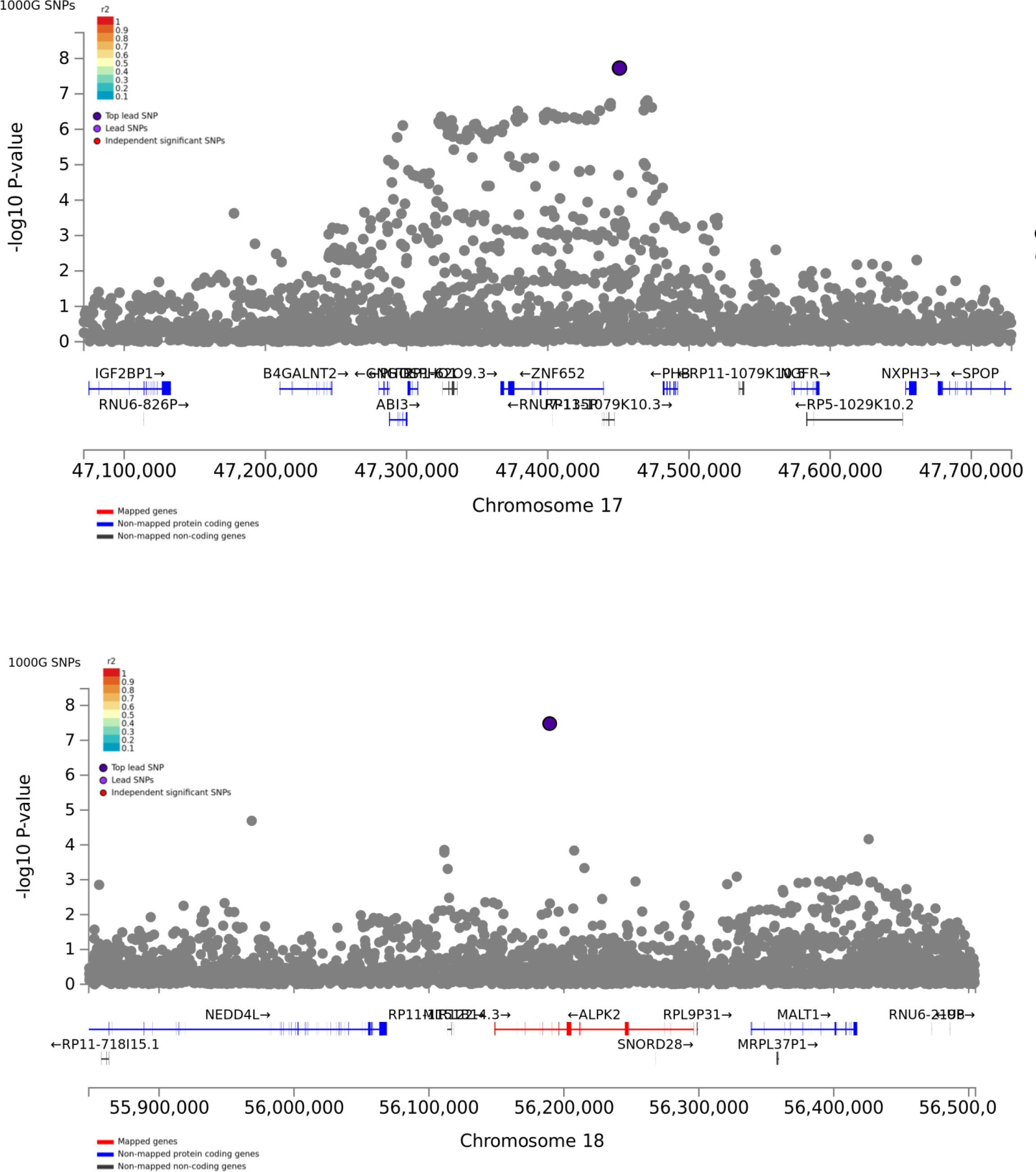
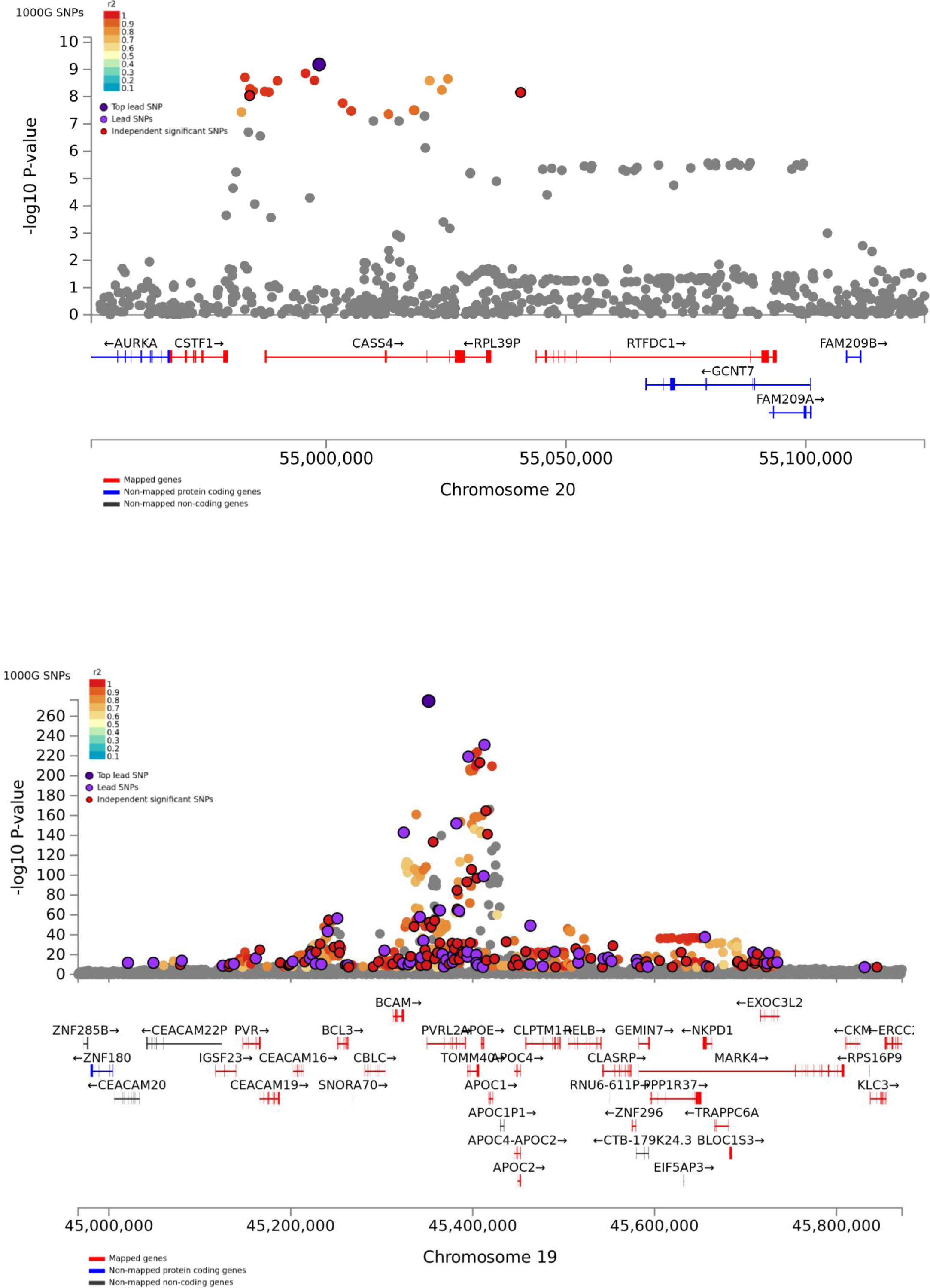
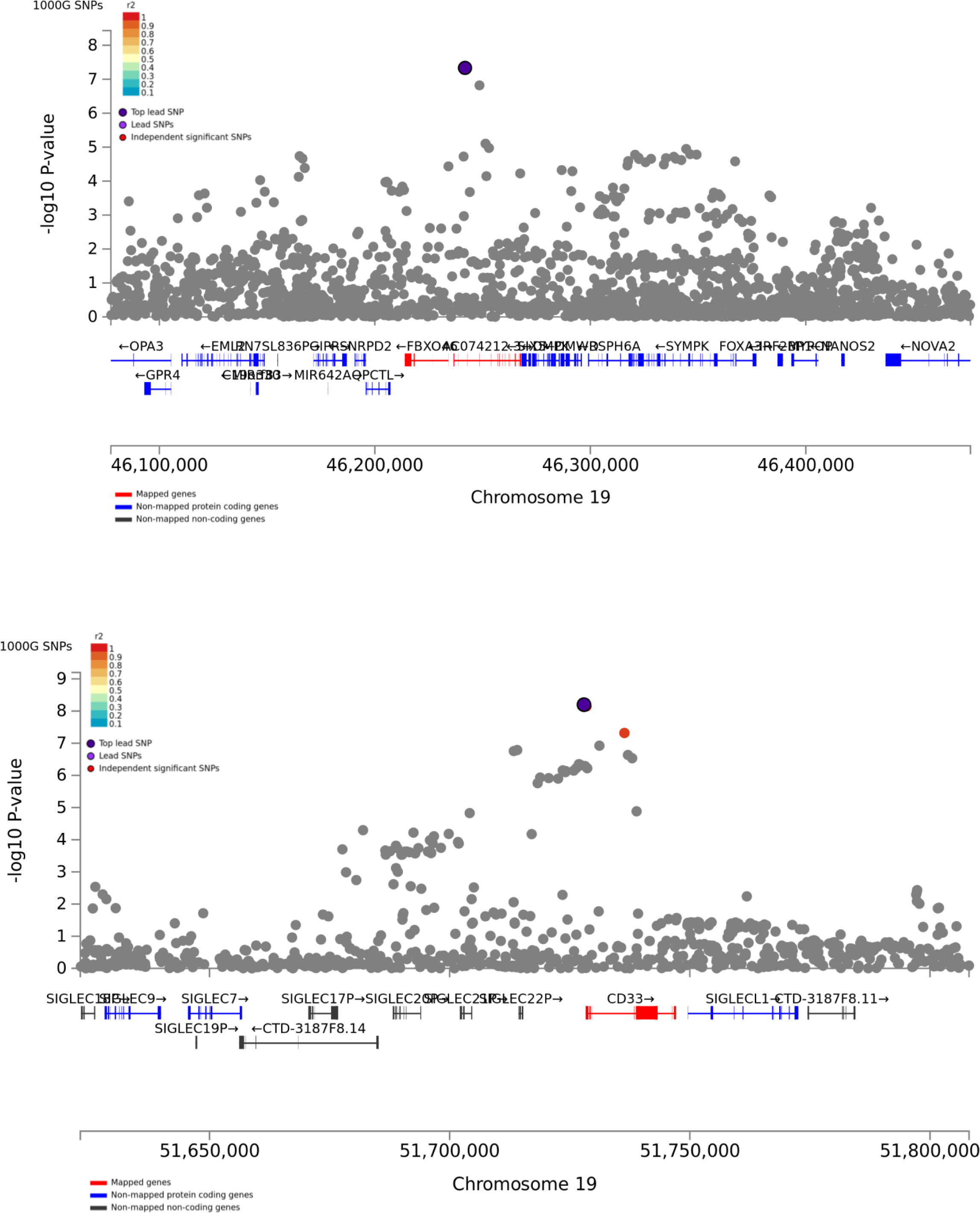
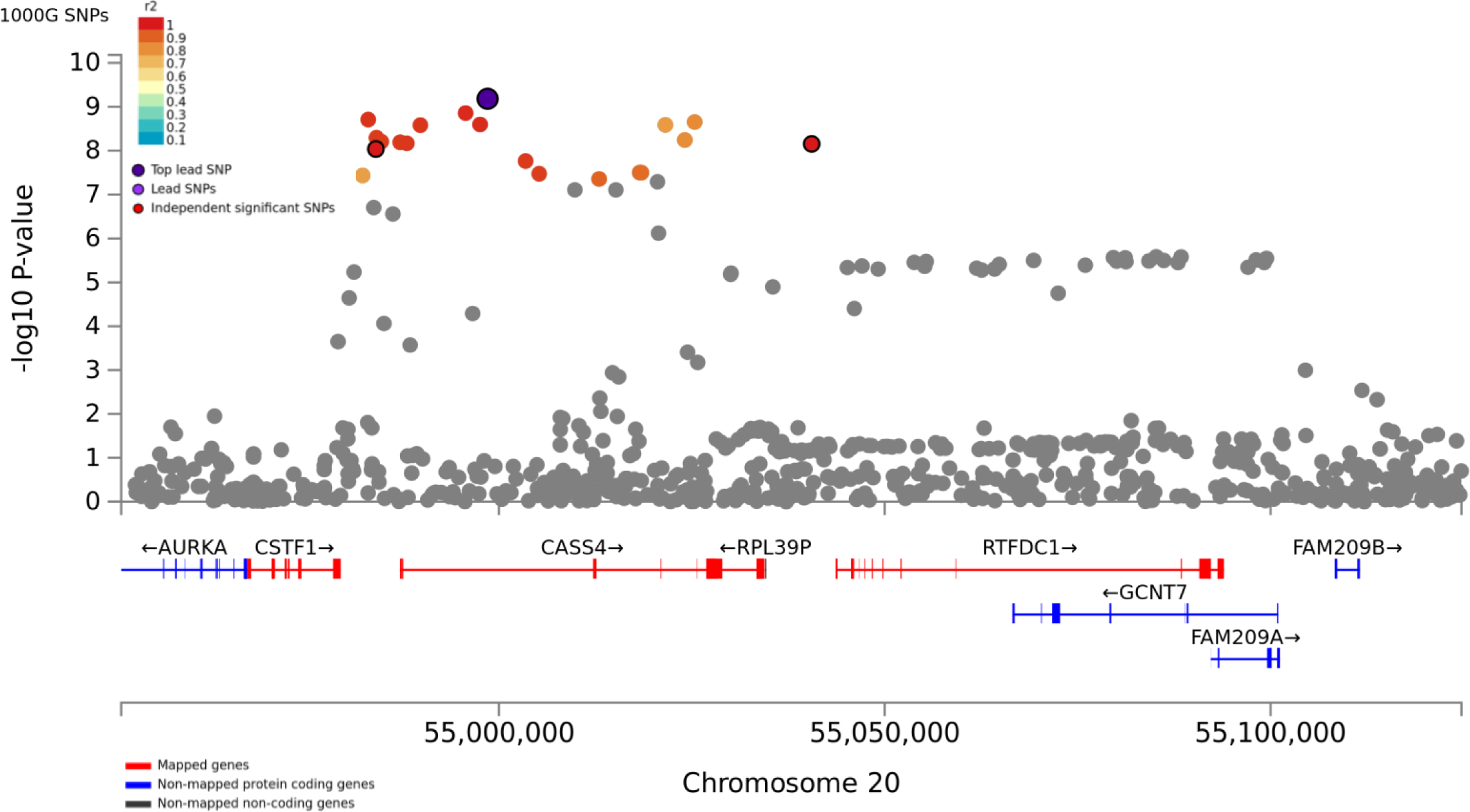
Regional plot for the 29 significant loci of the meta-analysis. Every point represents a SNP, which are colour-coded based on the highest r2 to one of the most significant SNPs, if greater or equal to r2 of 0.6. Other SNPs are coloured in grey.

**Supplementary Figure 3.**
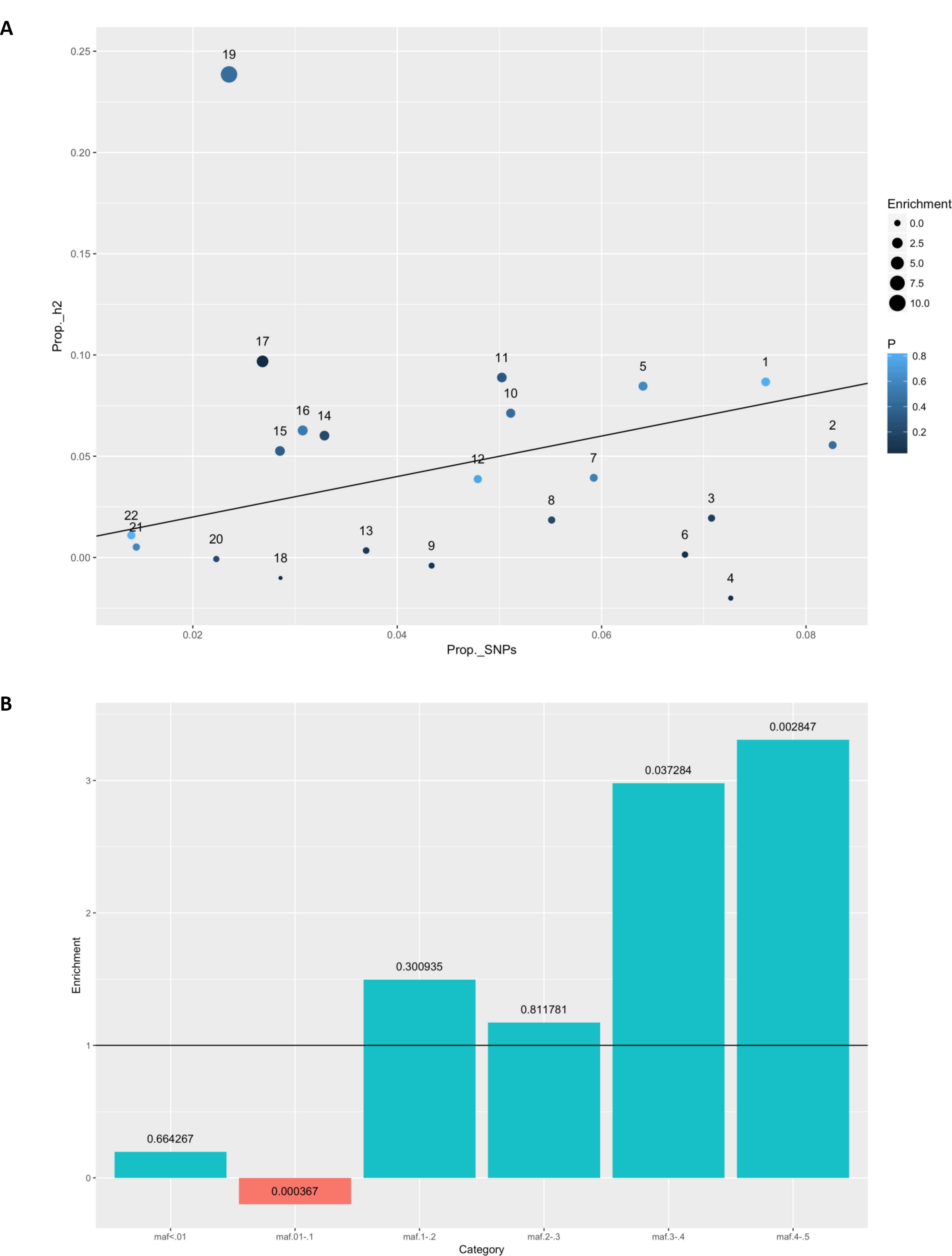
Partitioned heritability results for the meta-analysis. Variants were binned by chromosome or minor allele frequency and tested for a significant over or underrepresentation as to what is expected by chance. A) Enrichment results for heritability calculations where variants have been partitioned per chromosome. B) Enrichment results for heritability calculations where variants have been partitioned into multiple categories based on minor allele frequency.

**Supplementary Figure 4.**
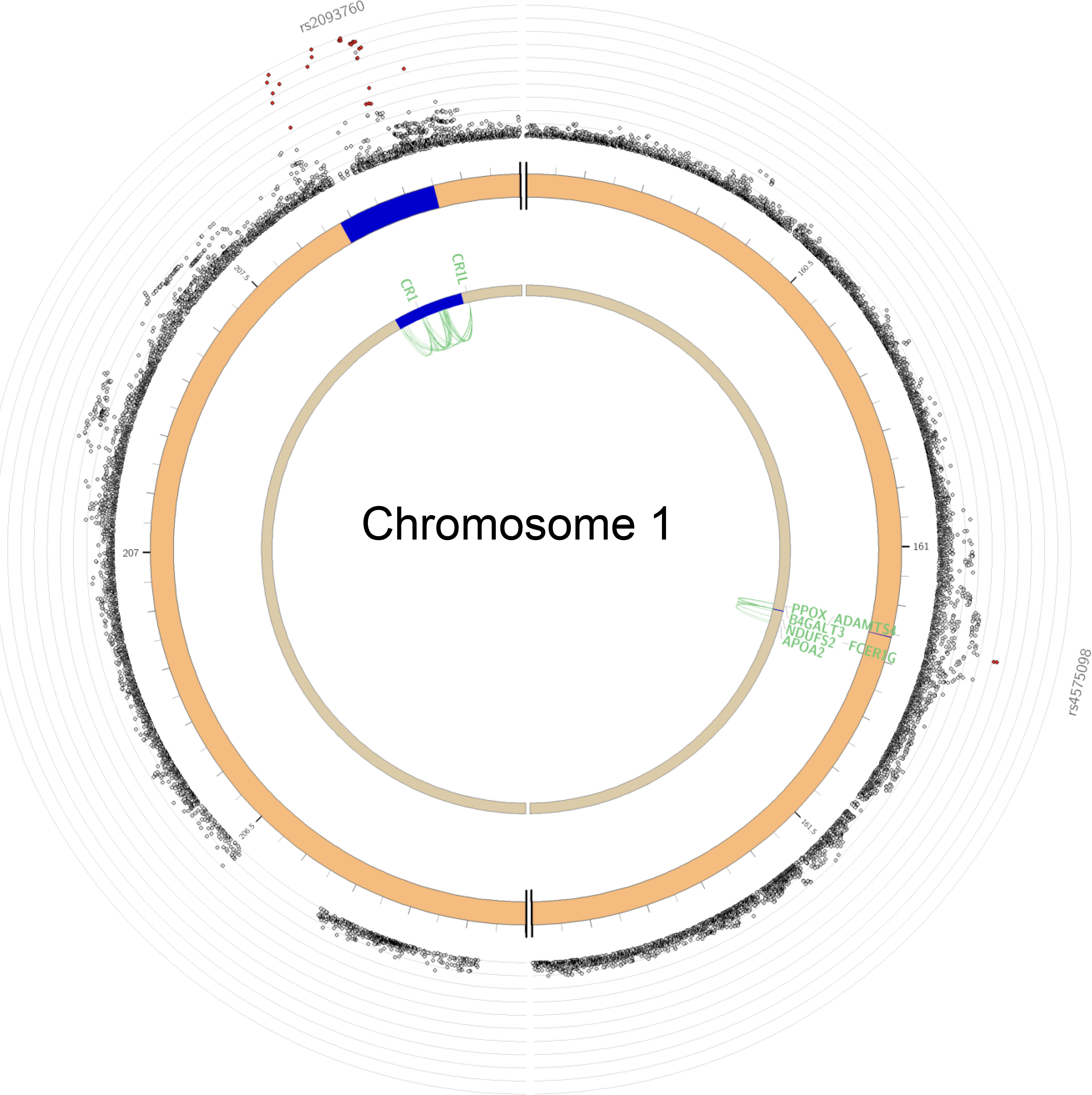
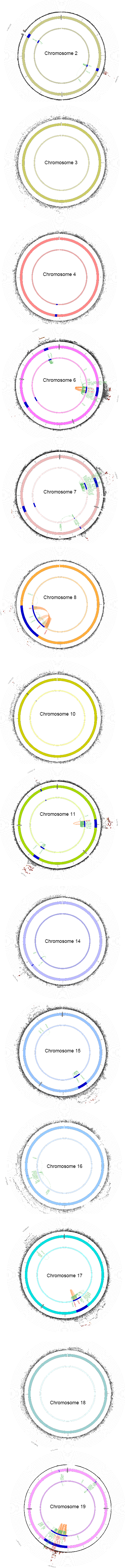
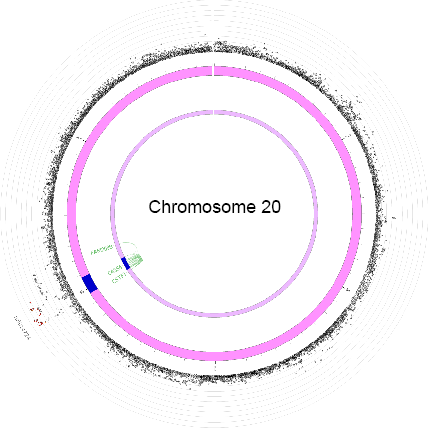
Full circos plots of chromatin interactions and eQTLs for all chromosomes with significantly associated loci. The distinct layers and colors correspond to various features. The outer layer contains zoomed in Manhattan plots containing only SNPs with P < 0.05. SNPs in genomic risk loci are color-coded as a function of their maximum r2 to the one of the independent significant SNPs in the locus, as follows: red (r2 > 0.8), orange (r2 > 0.6), green (r2 > 0.4) and blue (r2 > 0.2). SNPs that are not in LD with any of the independent significant SNPs (with r2 ≤ 0.2) are grey. The second layer displays the position of the genomic risk loci in blue. The third layer contains the mapped genes that are implicated by chromatin interactions and/or eQTL analysis (orange = chromatin interaction; green = eQTL; red = chromatin interaction and eQTL).

**Supplementary Figure 5.**
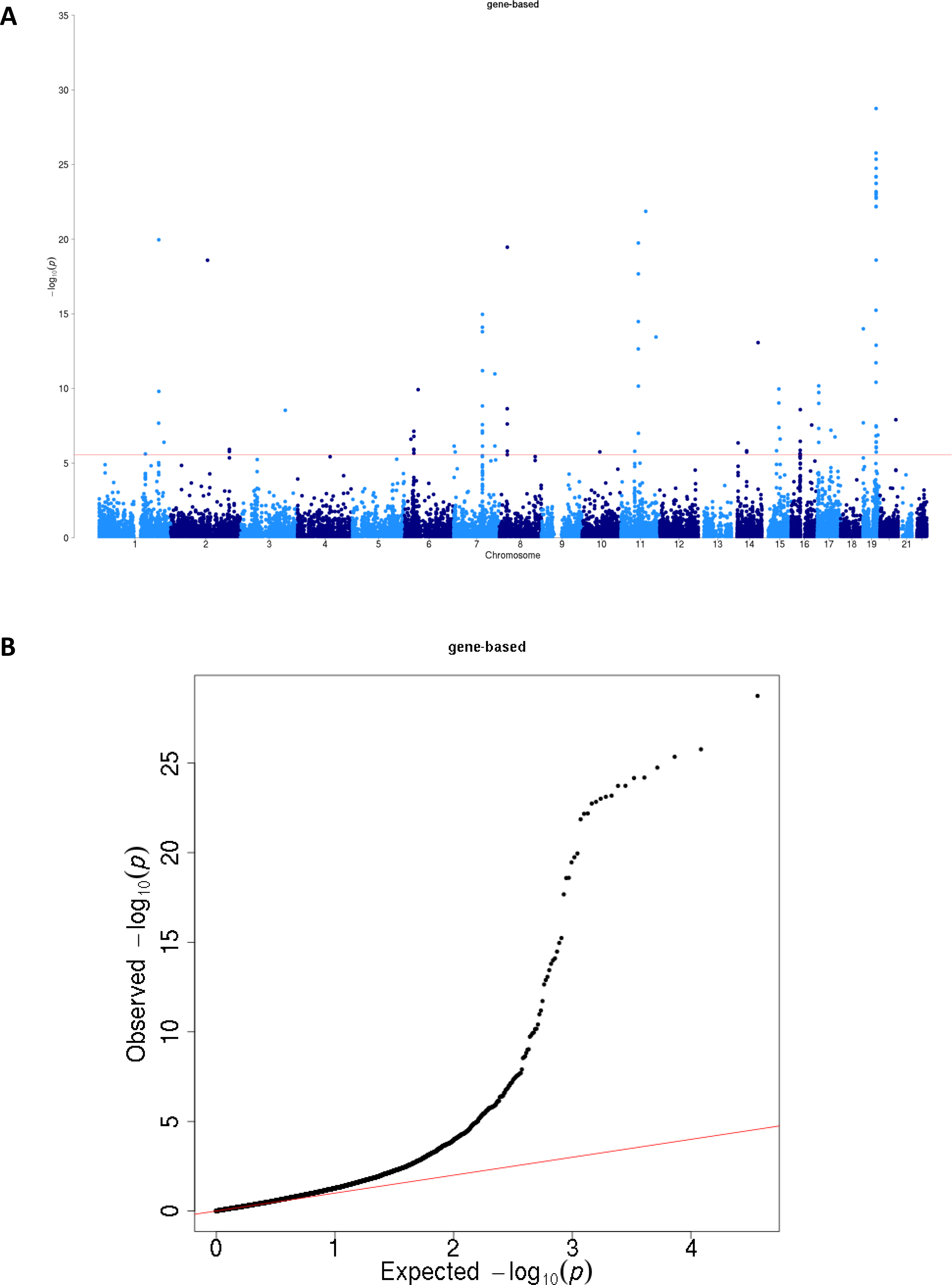
Gene-based association results with MAGMA. A) The Manhattan plot displays all associations per gene ordered according to their genomic position (start of gene) on the x-axis and showing the strength of the association with the −log10 transformed P-values on the y axis. B) The QQ plot displays the expected −log10 transformed p-values on the x-axis and the observed −log10 transformed p-values on the y-axis.

**Supplementary Figure 6.**
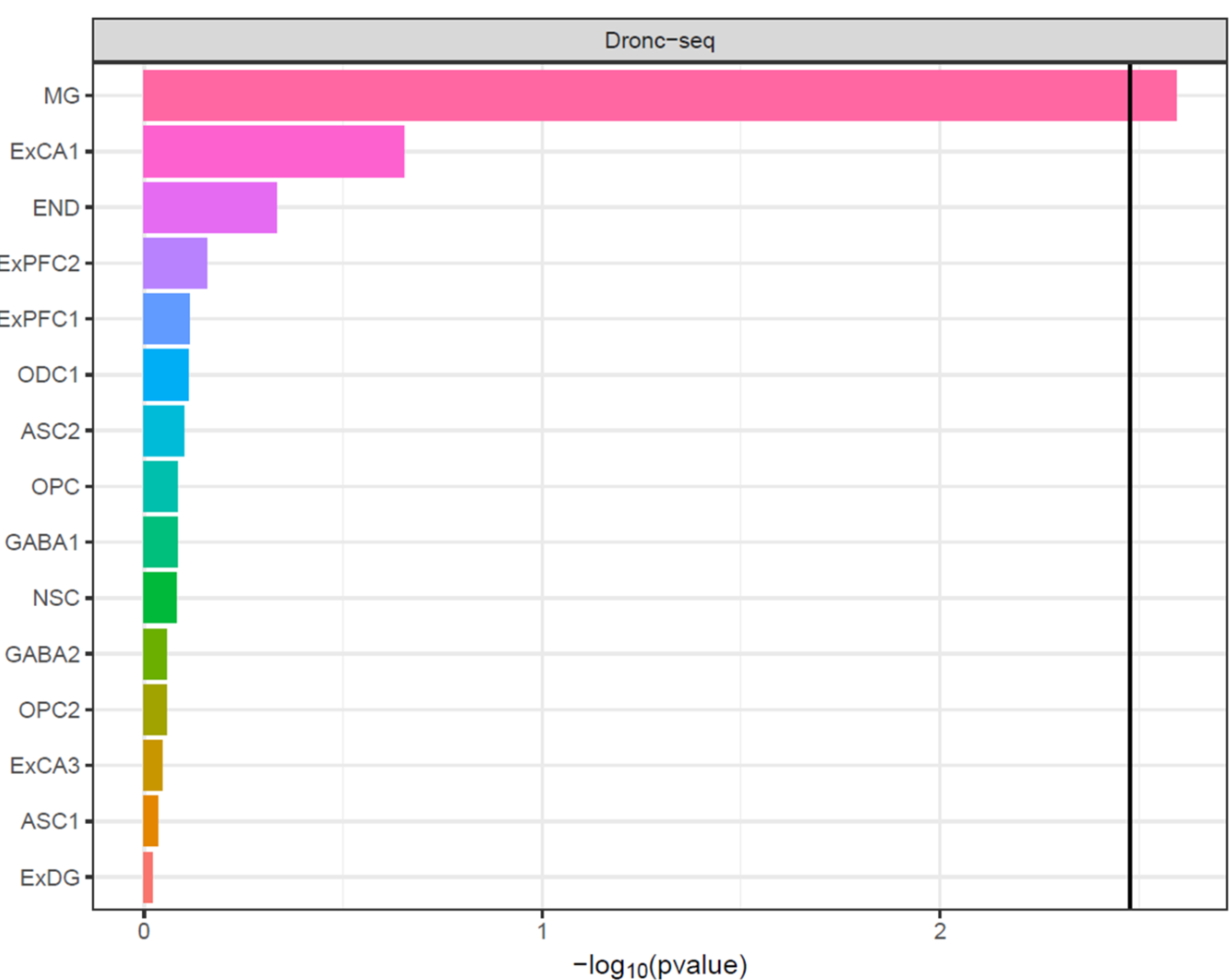
Single-cell expression gene-set results of human brain tissue. The black vertical line indicates the significance threshold correcting for number of tests within category. MG = microglia; ExCA1 = Hippocampal CA 1 pyramidal neurons; END = Endothelial cells; ExPFC2 = Prefrontal glutamergic neurons 2; ExPFC1 = Prefrontal glutamergic neurons 1; ODC1 = Oligodendrocytes; ASC2 = Astrocytes 2; OPC = Oligodendrocyte precursor cells 1; GABA1 = GABAergic interneurons 1; NSC = Neuronal stem cells; GABA2 = GABAergic interneurons 2; OPC2 = Oligodendrocyte precursor cells 2; ExCA3 = Hippocampal CA 3 pyramidal neurons; ASC1 = Astrocytes 1; ExDG = Dentate gyrus granule neurons.

**Supplementary Figure 7.**
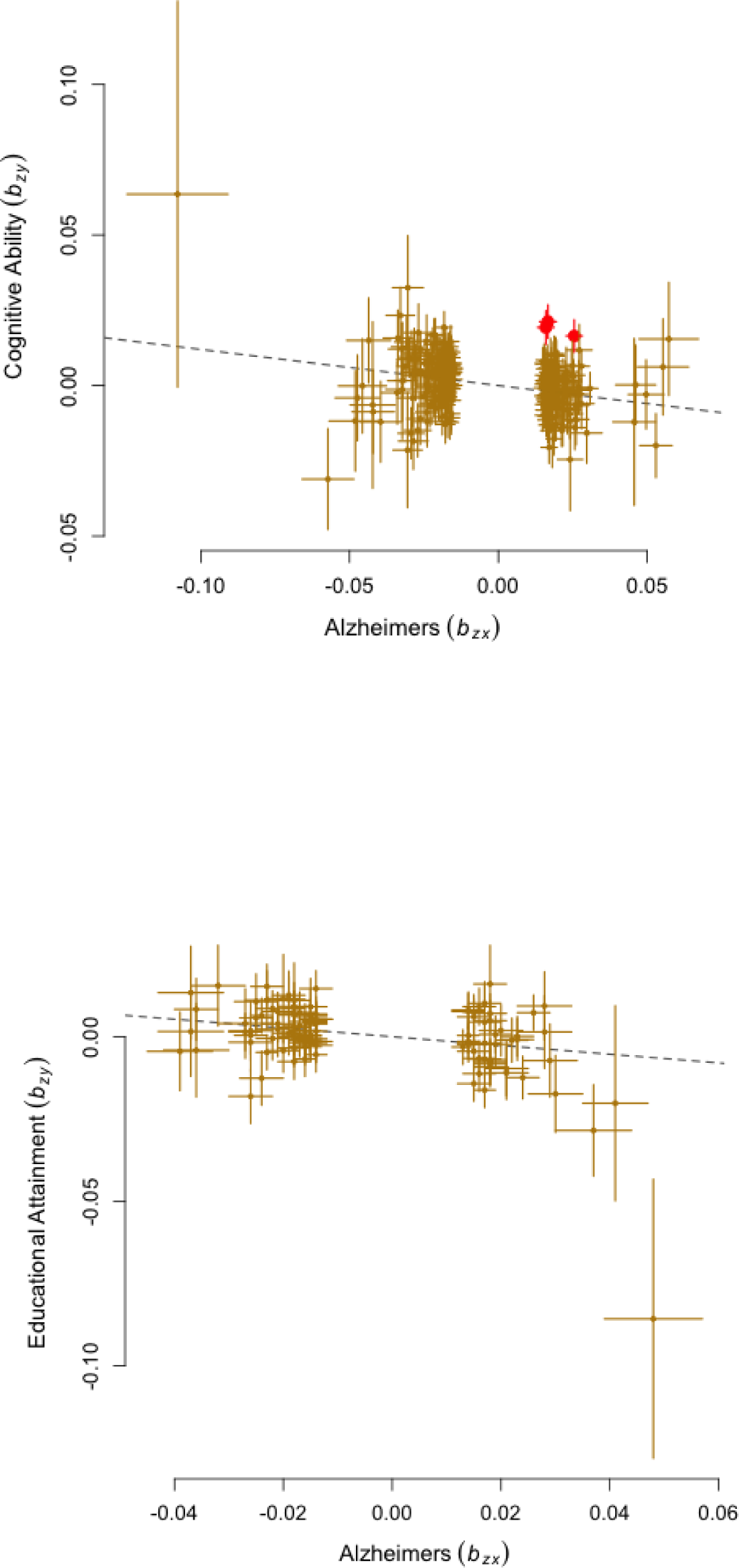
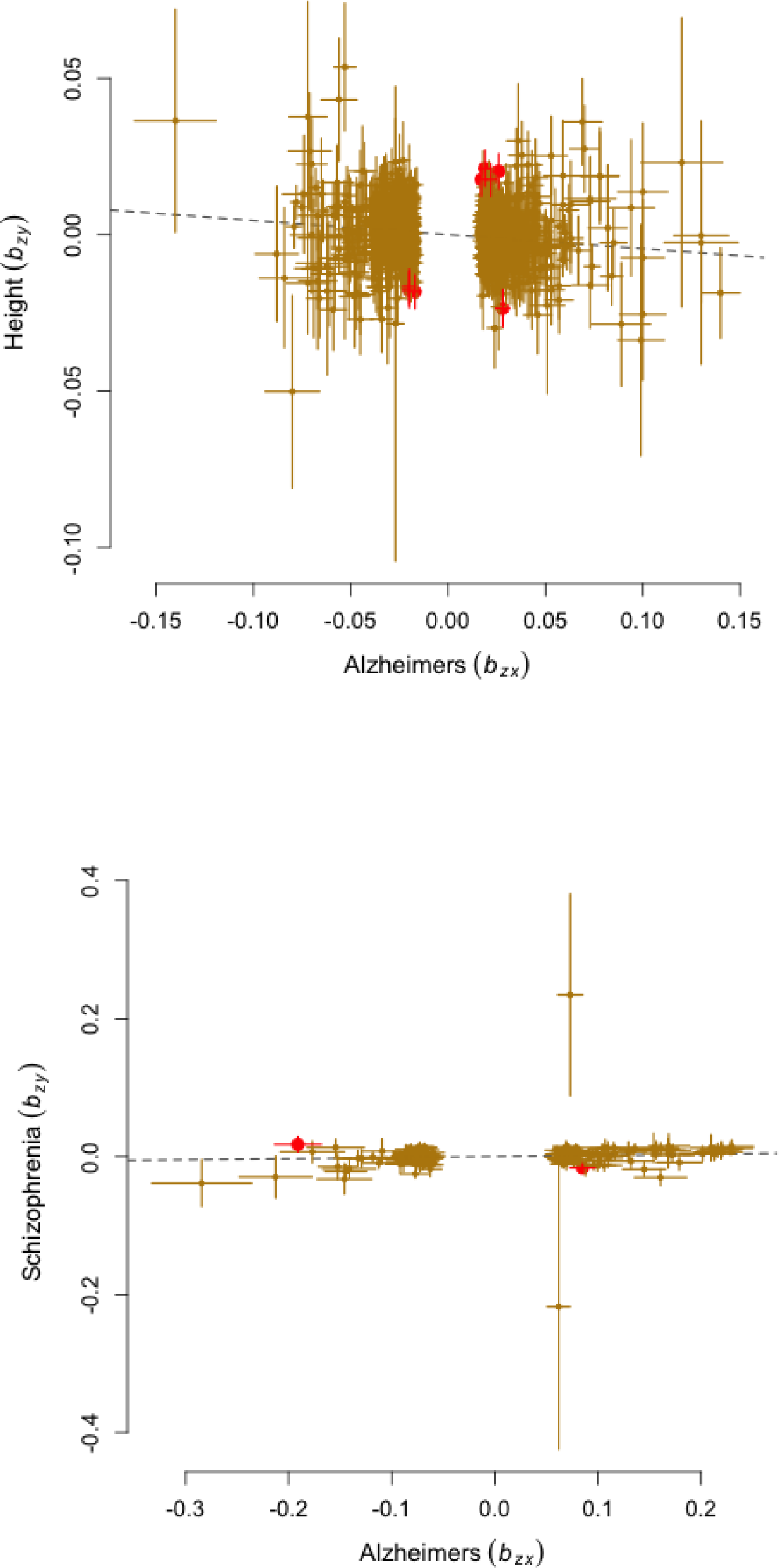
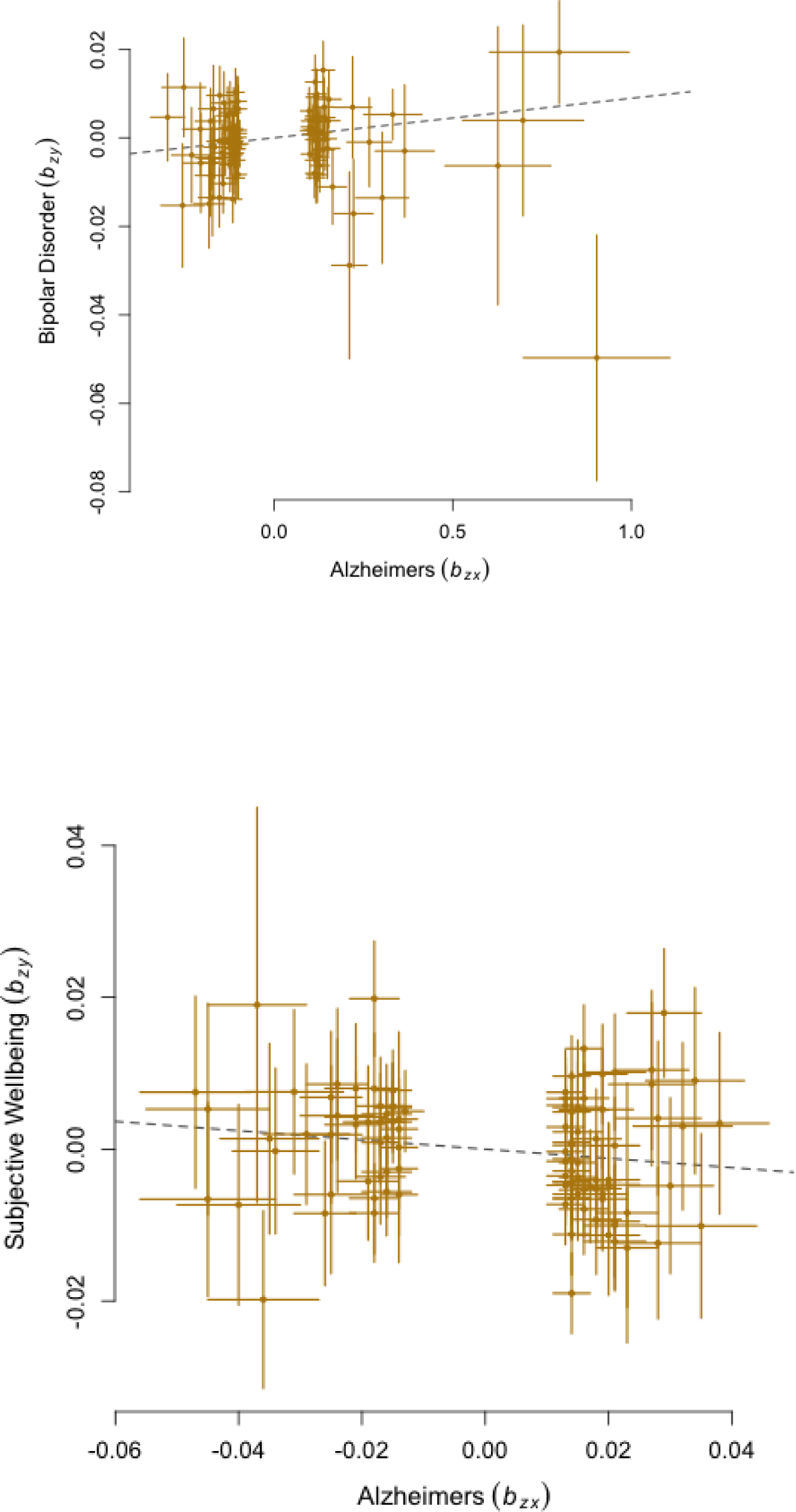
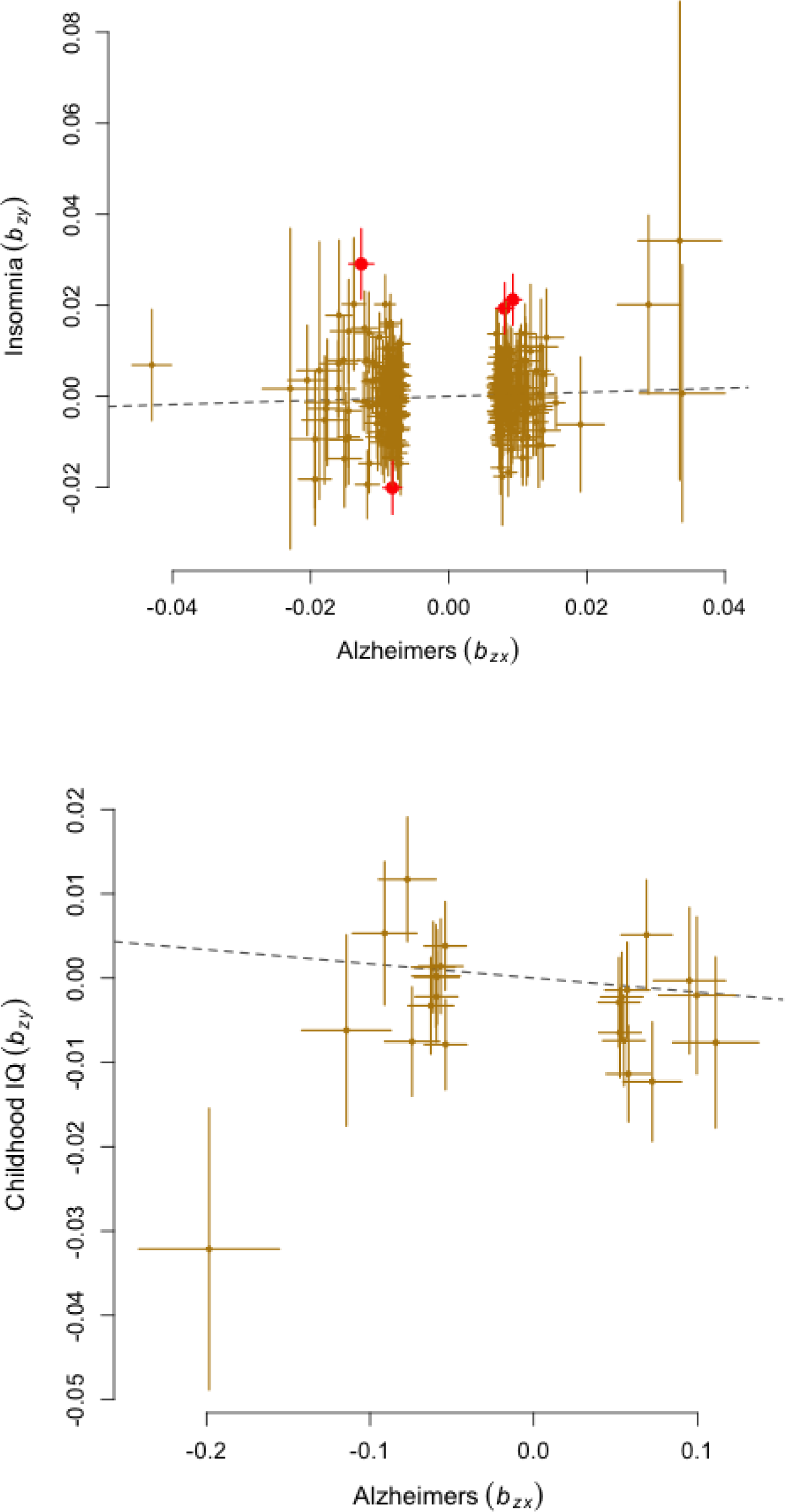
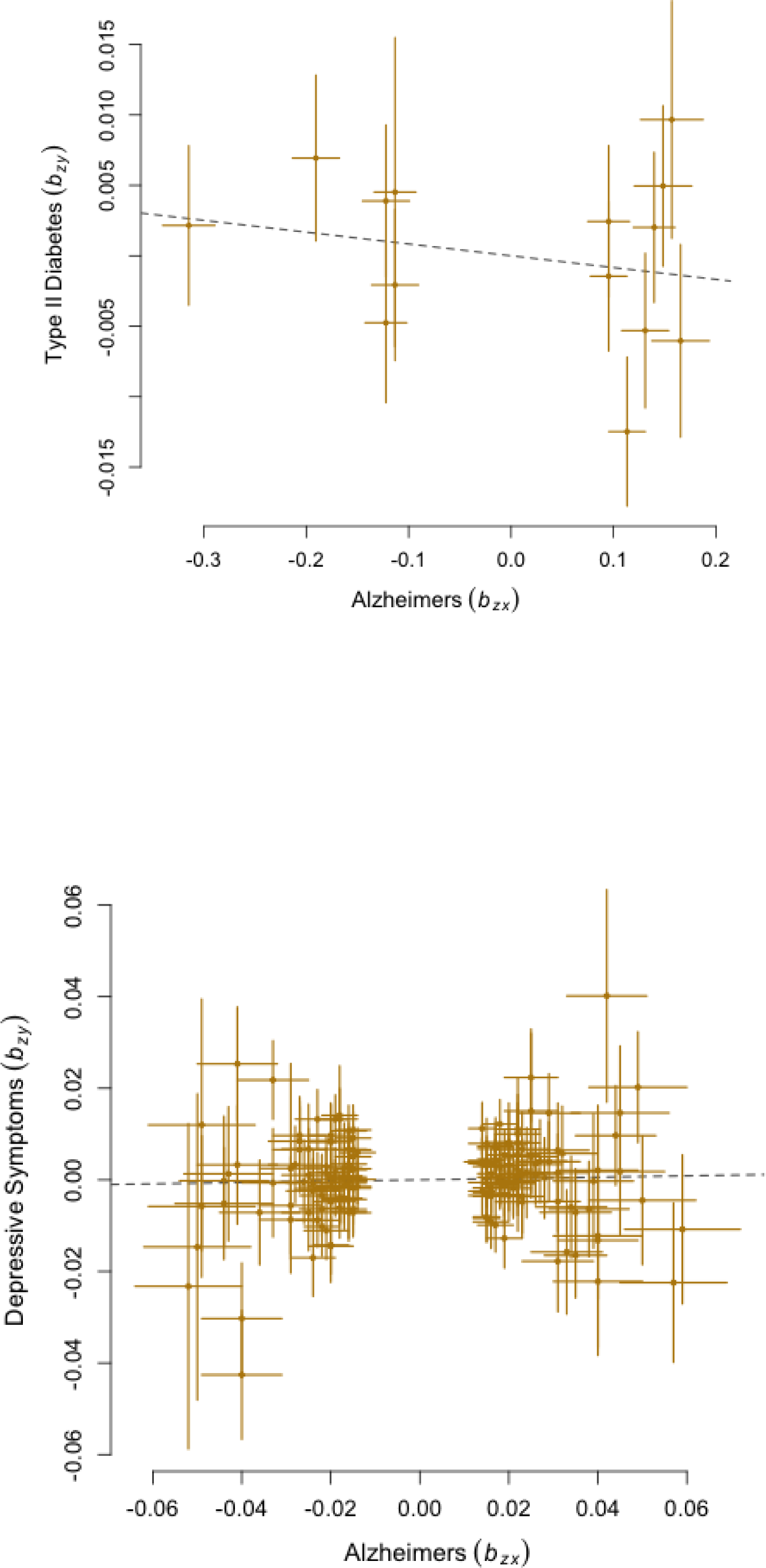
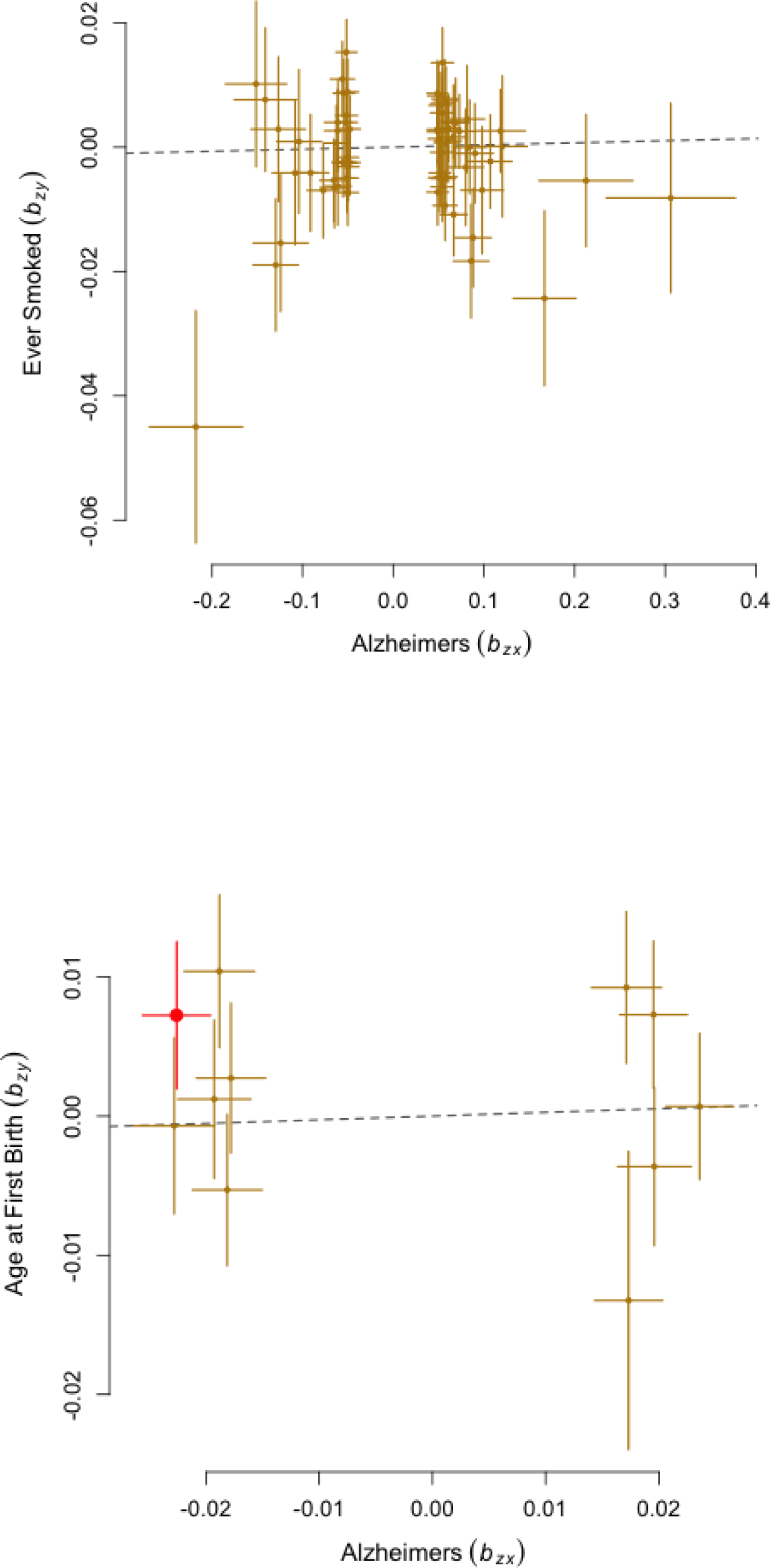
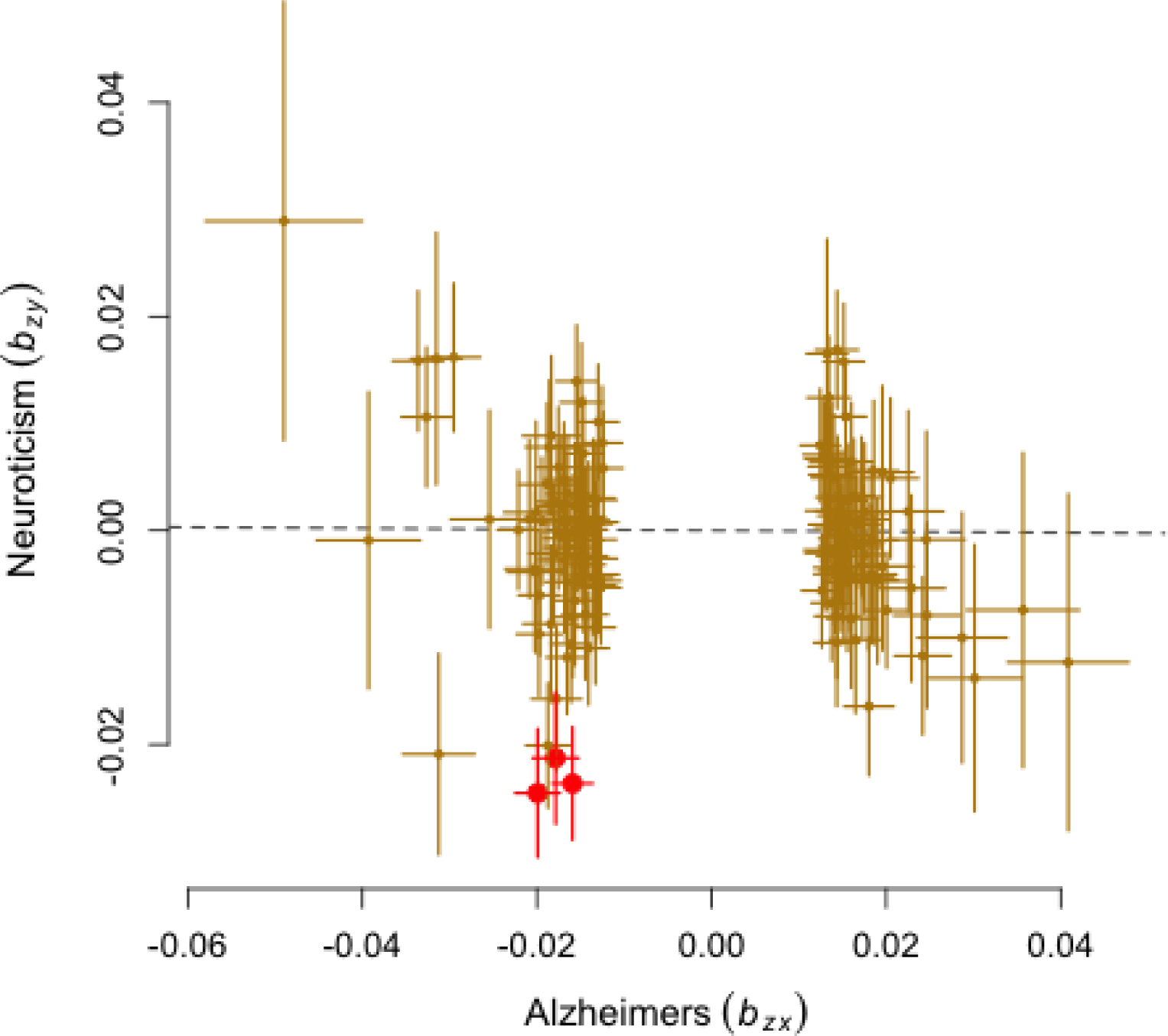
Mendelian Randomization tests for the effect of correlated phenotypes on risk for Alzheimer′s disease. For independent significant SNPs from each correlated phenotype, effect sizes of the SNPs for Alzheimer′s disease (**b_zy_**) are shown on the x-axis and effect sizes for correlated phenotypes are on the y-axis (b_zx_). The dotted line represents a line with slope of (*b_xy_*) and an intercept of 0.Red dots represent outliers that were excluded for the Mendelian Randomization analysis.

## Footnotes

* Although in other AD-related manuscripts this is common, we choose not to report the gene that is in closest proximity to the lead SNP as the ID for the locus, as this incorrectly implies that the gene is the causal gene for AD pathogenesis. We therefore believe it is preferred to use the rs-number of the most strongly associated SNP as an ID for the locus, and aim to highlight the most likely causal genes with more sophisticated functional interpretation analyses in later sections of this study.

* For straightforward comparison to this GWAS, we do here report the genes in closest proximity to the lead SNP. However, we would like to point out that GWAS findings implicate a genomic locus, and that the closest gene is not necessarily the causal gene.

